# The somatic mutation landscape of the human body

**DOI:** 10.1101/668624

**Authors:** Pablo E. García-Nieto, Ashby J. Morrison, Hunter B. Fraser

## Abstract

Somatic mutations in healthy tissues contribute to aging, neurodegeneration, and cancer initiation, yet remain largely uncharacterized. To gain a better understanding of their distribution and functional impacts, we leveraged the genomic information contained in the transcriptome to uniformly call somatic mutations from over 7,500 tissue samples, representing 36 distinct tissues. This catalog, containing over 280,000 mutations, revealed a wide diversity of tissuespecific mutation profiles associated with gene expression levels and chromatin states. We found pervasive negative selection acting on missense and nonsense mutations, except for mutations previously observed in cancer samples, which were under positive selection and were highly enriched in many healthy tissues. These findings reveal fundamental patterns of tissue-specific somatic evolution and shed light on aging and the earliest stages of tumorigenesis.

## INTRODUCTION

In humans, somatic mutations play a key role in senescence and tumorigenesis^1^. Pioneering work on somatic evolution in cancer has led to the characterization of cancer driver genes^2^ and mutation signatures^3^; the interplay between chromatin, nuclear architecture, carcinogens and the mutational landscape^4–7^; the evolutionary forces acting on somatic mutations^8–11^; and clinical implications of somatic mutations^12^.

Compared to cancer research, the study of somatic mutations in healthy tissues is limited. Early studies focused on blood^13,14^ as it is readily accessible and because of the known effects of immune-driven somatic mutation. Recently, somatic mutations have been characterized in tissues like skin^15^, brain^16,17^, esophagus^18,19^ and colon^20^. These studies confirmed that cells harboring certain mutations expand clonally, and the number of clonal populations—as well as the total number of somatic mutations—increases with age. Additionally, recurrent positively-selected mutations in specific genes (e.g. *NOTCH1*) were observed. However, a more comprehensive understanding of somatic mutations across the human body has been limited by the small number of tissues studied to date.

Most studies on somatic evolution in healthy tissues have sequenced DNA from biopsies to high coverage. However, the transcriptome also carries all the genomic information of a cell’s transcribed genome, in addition to RNA-specific mutations or edits. RNA-seq has been used to identify germline DNA variants^21^, and recently single-cell (sc) RNA-seq was used to call DNA somatic mutations in the pancreas of several people^22^.

To systematically identify somatic mutations in the human body and to investigate their distribution and functional impact, we developed a method that leverages the genomic information carried by RNA to identify DNA somatic mutations while avoiding most sources of false positives. We applied it to infer somatic mutations across 36 non-cancerous tissues, allowing us to explore the landscape of somatic mutations throughout the human body.

## RESULTS

### Somatic mutation calling across 36 non-disease tissues from 547 people

Our method considers genomic positions where we observed two alleles in the RNA-seq reads and assesses whether they are likely to be *bona fide* DNA somatic mutations (Fig 1a). We optimized the minimum levels of RNA-seq read depth, sequence quality, and number of reads supporting the variant allele to limit the impact of sequencing errors (Supp. Fig. 1a; see Methods). We then applied extensive filters to eliminate false-positives from biological and technical sources, including RNA editing, sequencing errors, and mapping errors (see Methods; Fig 1b-c, Supp. Fig. 2a; Supp. Table 1).

**Figure 1.**
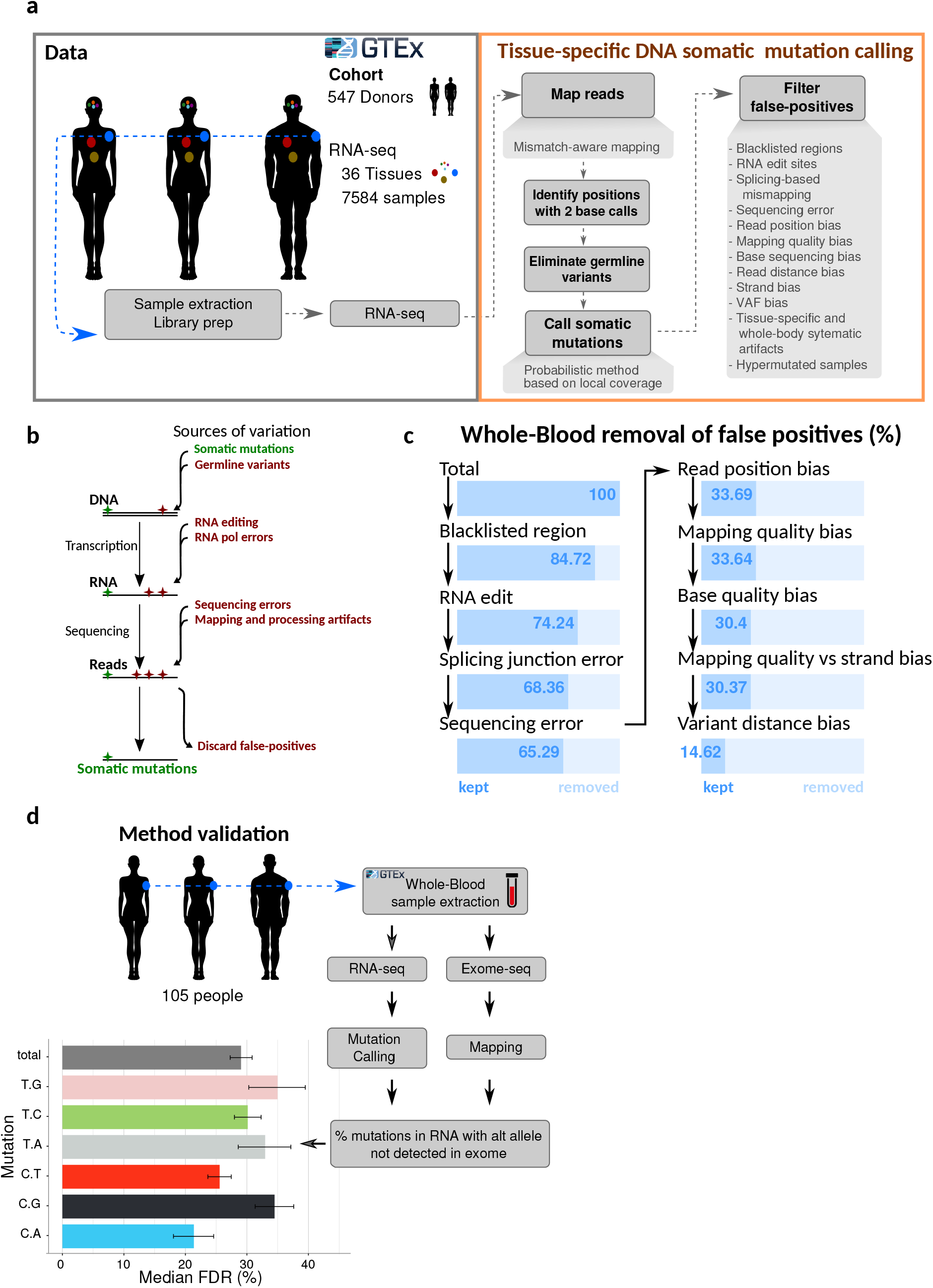
A method to identify DNA somatic mutations from RNA-seq. **a,** A general overview of the method. RNA-seq reads were downloaded from GTEx v7 (left) and processed to identify positions with two different base calls at a high confidence. Then, sources of biological and technical artifacts were removed (right, see Methods). **b,** Schematic illustrating potential sources of sequence variation. **c,** Average percentage of variants detected in blood RNA-seq that are retained after each step of filtering (see Methods). **d,** Validation of the method. For 105 individuals we compared variant calls from exome DNA-seq data with those from RNA-seq of the same samples. Median FDR values per mutation type are shown and they represent the fraction of mutations called in RNA-seq for which there are no exome reads supporting the same variant (see Methods and Supplementary Figure 1c). Error bars represent the 95% confidence interval after bootstrapping 10,000 times.

To validate the method, we compared somatic mutation calls from 105 blood RNA-seq samples to exome DNA sequencing performed on the same samples^23^ (Fig. 1d). We observed a false-discovery rate (FDR) of 29% which represents the percentage of somatic mutations called from RNA-seq not having evidence in the corresponding DNA exome-seq sample (Fig. 1d, Supp. Fig. 1c; see methods). This is comparable to the 40% FDR in a previous study that inferred mutations from scRNA-seq in pancreas^22^.

After applying the pipeline and filters to RNA-seq data from the GTEx project, we retained a total of 7,584 samples from 36 different tissues and 547 different individuals with no detectable cancer (Supp. Table 2). This resulted in a total of 280,843 unique mutations (Supp. Table 3), most of which were rare across the entire data set (median frequency = 0.026% of samples; Supp. Fig. 2b).

We first investigated the factors influencing mutation counts per sample and tissue (see Methods). The main contributor was sequencing depth and to a lesser extent other biological and technical factors (Supp. Fig. 2c, Supp. Table 4). Tissues that have more mutations than expected from sequencing depth include those most often exposed to environmental mutagens or with a high cellular turnover like skin, lung, blood, esophagus mucosa, spleen, liver and small intestine (Fig. 2a). On the other end of the spectrum are those with low environmental exposure or low cellular turnover such as brain, adrenal gland, prostate, and several types of muscle — heart, esophagus muscularis, and skeletal muscle (Fig. 2a).

**Figure 2.**
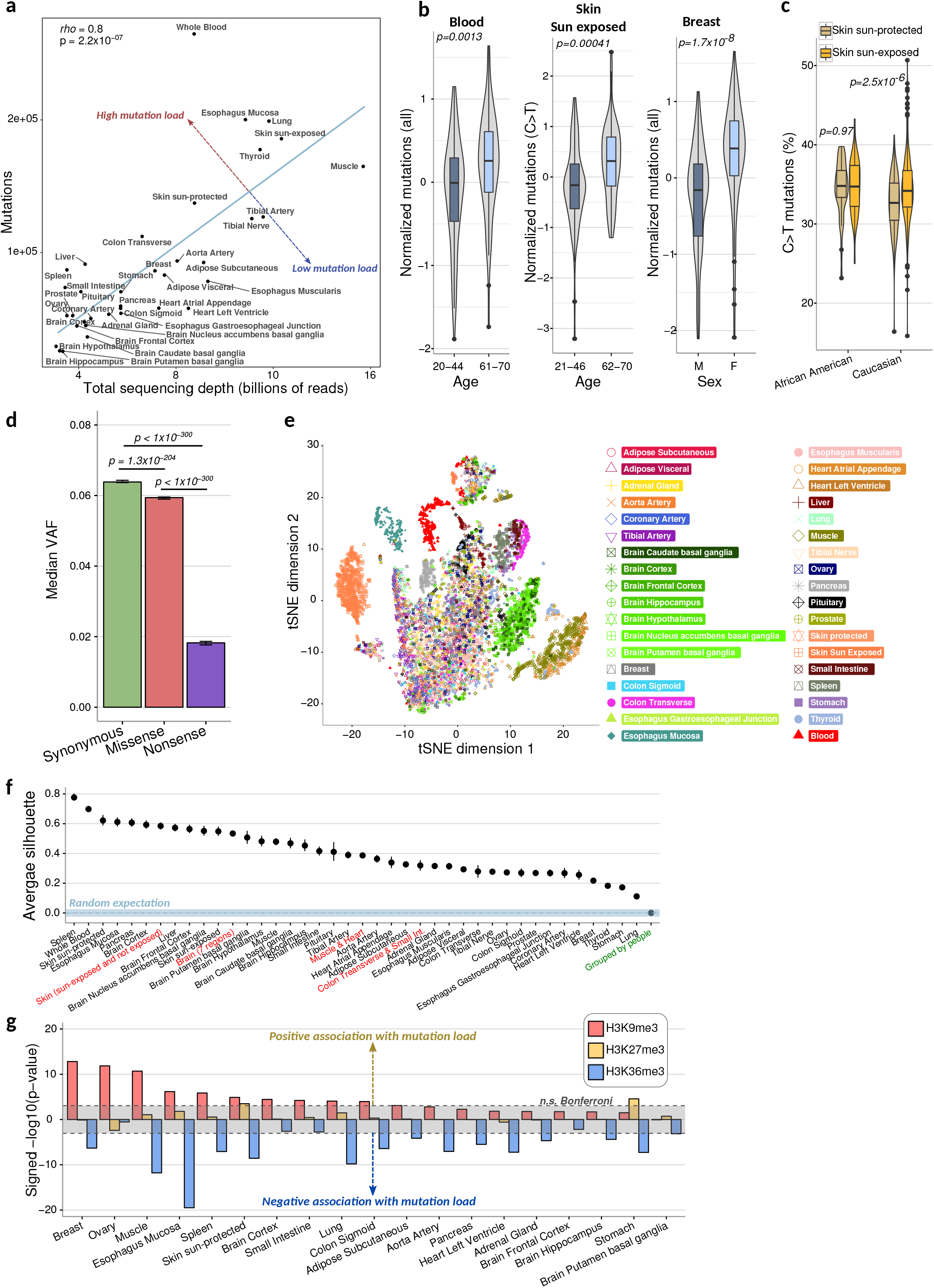
Cross-tissue analysis of somatic mutations. **a,** The total number of mutations observed in a tissue depends on the sequencing depth of that tissue. Sequencing depth is defined as the cumulative amount of uniquely mapped reads across all samples of a tissue. A linear regression line is shown in blue; tissues above it exhibit more mutations than expected by sequencing depth, while tissues below it show fewer mutations than expected. rho is the Spearman coefficient. **b,** Examples of significant mutation associations with age and biological sex (see Supplementary Figure 4 and Supp. Table 6 for all tissue data). Age ranges represent the youngest and oldest quartiles for each tissue. To control for sequencing depth and other technical artifacts, mutation values were obtained as the residuals from a linear regression (see Methods). P-values are from a two-sided Mann-Whitney test. **c,** Caucasian sun-exposed skin shows a higher percentage of C>T mutations compared to sun-protected skin, while no such difference was seen for African-American skin. P-values are from two-sided Mann-Whitney tests. **d,** Median variant allele frequency (VAF) for each mutation type based on their impact to the amino acid sequence; error bars represent the 95% confidence interval after bootstrapping 1000 times; p-values are from two-sided Mann-Whitney tests. **e,** tSNE plot constructed from a normalized penta-nucleotide mutation profile (the mutated base plus two nucleotides in each direction; see Methods for normalization details) and all samples in this study. **f,** Average silhouette scores representing the coherence of selected groups of samples from the tSNE space in panel e; a score of 1 represents maximal clustering, whereas 0 represents no clustering (see Methods). Grouping was performed by tissue-of-origin, or multiple tissues combined (red labels). “Grouped by people” (green label) is an average silhouette score after grouping samples by their person-of-origin from 20 randomly selected people. The blue dashed line represents the average random score expectation after permuting tissue labels (see Methods) and the blue stripes are +/− two standard deviations. Error bars in points represent the 95% confidence interval based on bootstrapping 10,000 times. **g,** Mutation load is positively associated with H3K9me3 and/or negatively associated with H3K36me3 across most tissues analyzed. P-value of association was obtained from a linear regression using all chromatin marks as explanatory variables (see Methods). Gray range denotes non-significant p-values after Bonferroni correction.

We observed similar trends across all six possible point mutation types (complementary mutations – such as C>T and G>A – were collapsed because when one is detected the other is always present on the other DNA strand) (Supp. Fig. 3a). C>T and T>C mutations were the most abundant types followed by C>A (Supp. Fig. 3b; Supp. Table 5).

### Mutation load is affected by age, sex, ethnicity, and natural selection

After controlling for all known technical factors (see Methods), we observed that age was positively correlated with mutation load across most tissues (Supp. Fig. 4a; 29/36 tissues with Spearman *ρ* > 0, binomial p = 1.6×10^−4^). Blood showed the most significant age association for all mutation types combined (Fig. 2b; 1^st^-4^th^-quartile Wilcoxon p = 0.013; Spearman *ρ* = 0.21, Benjamini-Hochberg [BH] FDR < 0.0001), whereas sun-exposed skin was the most significant for C>T mutations, the most common mutation associated with UV radiation (1^st^-4^th^-quartile Wilcoxon p = 0.00041; Spearman *ρ* = 0.17 BH-FDR = 0.02). We observed other strong age associations in several brain regions, and in particular the basal ganglia (Supp. Fig. 4a), supporting the hypothesis that somatic mutation load could contribute to the increasing risk of neurodegenerative diseases with age^24^.

In addition to age, other biological factors also contributed to somatic mutations. For example, women show a much greater mutation load in breast than men (Fig 2b; Wilcoxon p = 1.7×10^−8^). We also observed female-biased mutations (BH-FDR<0.1) in subcutaneous adipose, visceral adipose, liver, and the adrenal gland; in contrast, we found no significant male-biased mutations after multiple hypothesis correction (Supp. Fig. 4c; Supp. Table 6). Ethnicity can also affect mutation rates: we found a significant increase of C>T mutations in Caucasian sun-exposed skin compared to non-exposed skin, but no corresponding difference in African-Americans (Fig. 2c, Supp. Fig. 4b), likely due to protection against UV radiation provided by higher melanin content. Finally, we observed that the number of stem cell divisions in a tissue was weakly correlated with mutation load (see Supp. Note 1, Supp. Fig. 5).

To investigate the role of natural selection on deleterious somatic mutations, we examined the variant allele frequency (VAF) within every sample where a given mutation was observed. If mutations are subject to negative selection, we would expect it be acting most weakly on synonymous mutations that do not change the amino acid sequence; more strongly on missense variants that change an amino acid; and most strongly on nonsense mutations that introduce a premature termination codon. Indeed, this was the pattern we observed, with synonymous variants having significantly higher VAF than both missense and nonsense variants (Fig. 2d).

Since our mutations are called from the transcriptome, these results could be influenced by Nonsense Mediated Decay (NMD), which degrades transcripts containing premature termination codons. While we did observe evidence of NMD (Supp. Fig. 4d), we found that nonsense mutations in the last exon of genes – which are not subjected to NMD^25^ – still have significantly lower VAF compared to synonymous and missense mutations (Wilcoxon p = 1.9×10^−213^ and p = 5.6×10^−150^, respectively; Supp. Fig. 4d), confirming the existence of negative selection against nonsense mutations. Therefore, in contrast to what has been observed in cancer^26^, we found evidence of negative selection acting against both missense and nonsense mutations.

### Somatic mutation profiles are tissue-specific and associated with chromatin state

To visualize the tissue-specificity of somatic mutational patterns, we applied t-SNE^27^ to the full set of mutations called in each of the 7584 tissue samples (including the two bp flanking each mutation; see Methods; Fig 2e). We then used a silhouette score (SS) to quantify clustering of samples in this two-dimensional space (see Methods), where a value *SS = 1* indicates samples that are maximally clustered within a group, and a value *SS=0* means that samples are equidistant to samples within vs. outside a group. Grouping samples by their donor-of-origin results in a mean *SS = 0*, no different than expected by chance (Fig. 2f). However, grouping samples by tissue resulted in values *0.7 > SS > 0.1*, suggesting tissue-specific mutation patterns. Spleen, blood, skin, liver, and esophagus mucosa exhibited the most coherent profiles (Fig. 2f, Supp. Fig. 6a,d); conversely artery, lung, stomach and thyroid had the least consistent profiles (Fig. 2f, Supp. Fig. 6e,f). Interestingly, some tissues cluster together as shown by SSs obtained by grouping samples from more than one tissue, suggesting shared mutagenic or repair processes (Fig 2f; Supp. Fig. 6b,c; e.g. skeletal muscle and heart, *SS = 0.38*; seven brain regions, *SS = 0.53*; colon and small intestine, *SS = 0.31*). These results suggest that the somatic mutation landscape of the transcribed genome is largely defined by tissue-of-origin.

To explore the source of this tissue-specificity, we hypothesized that chromatin may play an important role, as it does in cancer^4,5^. We assessed the association between tissue-specific mutation rates and five chromatin marks measured across many human tissues by the Roadmap Epigenomics Project^28^ (Supp. Table 7, see Methods). Across most tissues (except brain) there is a strong positive association between mutation rate and a marker for heterochromatin (H3K9me3); conversely, we found strong negative associations with actively transcribed chromatin (H3K36me3; Fig. 2g). These results suggest that chromatin associations with mutation rates in the human body are nearly ubiquitous and arise prior to cancer development.

### Mutational strand asymmetries are widespread and vary across individuals

Cancer mutations often occur preferentially on one DNA strand, either in reference to transcription or to DNA replication (leading vs lagging strand)^29^. Since our mutations are derived from transcribed exons, we focused on transcriptional mutational strand asymmetries.

We observed the strongest asymmetry for C>A mutations, which preferentially occur on the transcribed strand in most tissues except for brain (Fig. 3a,b; mean [transcribed/non-transcribed] ratio = 1.6 in non-brain, 0.98 in brain). C>A mutation load on the transcribed strand also showed the greatest variation between individuals (Fig. 3a,b). C>A asymmetries were often correlated between different tissues of the same person (Fig. 3c, Supp. Fig. 7a), suggesting a common factor can induce an over-accumulation of C>A mutations on the transcribed strand (or equivalently, G>T mutations on the non-transcribed strand) across many of an individual’s tissues. This factor is of unknown origin; it could be intrinsic (e.g. genetic), extrinsic (e.g. exposure to a mutagen), or a combination of the two. Interestingly, this same C>A asymmetry has been observed in lung and ovarian cancer^29^ and in cell lines exposed to different environmental agents^30^; our results suggest it is far more widespread, occurring in most tissues except for brain.

**Figure 3.**
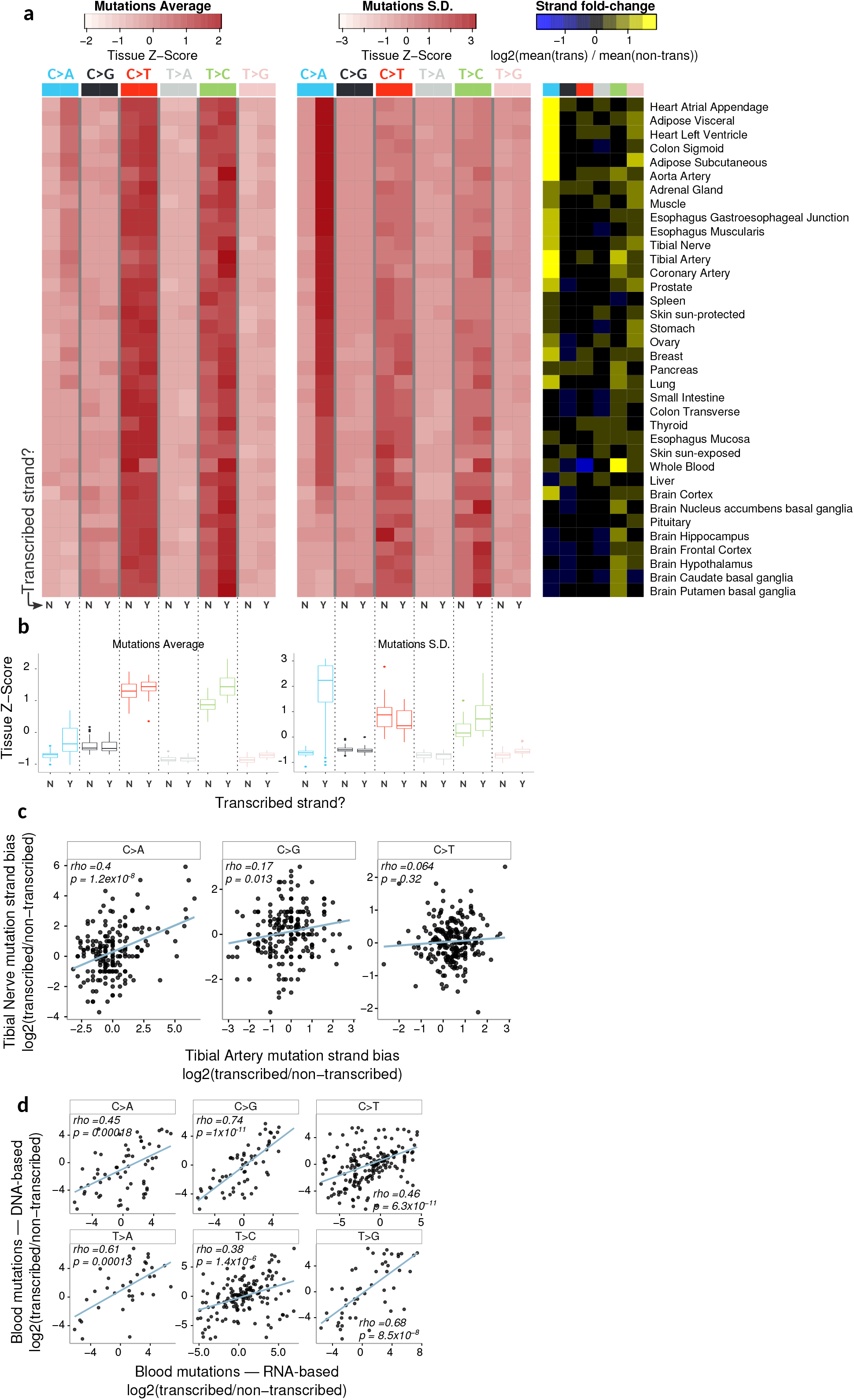
Somatic mutational strand asymmetries. **a,** Mutation average and S.D. for each strand with respect to transcription (left and middle panels) and the ratio of mean mutations on the transcribed over the non-transcribed strand (right panel). **b,** Distribution of z-scores for mutation averages and standard deviations on each strand (from panel a) across all samples. **c,** Example of intraindividual correlation of C>A strand asymmetry in two tissues; each point represents an individual for which we generated mutation maps in the two tissues (see Supp. Fig. 7a for all tissue pairwise comparisons). **d,** Correlation of mutational strand biases between calls from RNA-seq and matched DNA-seq; each point represents the mutation strand bias observed in one gene across all 105 blood samples. Blue lines in all scatter plots are based on a linear regression; rho values are the Spearman correlation coefficients.

Strand asymmetries could potentially arise from RNA-specific edits or transcriptional errors. To test if these are *bona fide* DNA mutations, we examined exome data from matching blood samples, and observed general agreement between the level of asymmetry per gene as measured by DNA vs. RNA-seq (Fig. 3d).

In addition, we found blood samples to have the highest levels for C>T and T>C asymmetries. However, the directionality of these asymmetries suggests that samples with high biases may be driven by the two major types of RNA editing: A>I (represented in our data by T>C on the transcribed strand), and C>U (represented in our data by C>T on the transcribed strand) (Fig. 3a, right panel). Interestingly, estimating cell type abundances in each blood sample (see Methods) revealed that the abundances of two cell types, resting NK cells and CD8+ T cells, showed the strongest associations with the extent of both of these asymmetries (Supp. Fig 7b,c). Consistent with this result, these same two cell types have been shown to have increased levels of both types of editing in stress conditions^31^. This suggests that some RNA editing sites may be present in our catalog of somatic mutations, though we did not observe an increased FDR for these two mutation types in blood (Fig. 1d), suggesting that RNA editing has not substantially inflated our FDRs.

### Gene expression implicates pathways associated with somatic mutation load

To explore cellular factors accompanying an increase in somatic mutation load across non-disease tissues, we performed an unbiased tissue-level search for genes whose expression was associated—either positively or negatively—with exome-wide mutational load (see Methods). These enrichments may reflect a mixture of causal scenarios: gene expression impacting mutations or vice versa, or both driven by a third variable. Here we focused on the most-abundant C>T mutations; other mutation types are described in the supplement (Supp. Figs. 9,10).

While most genes were tested in the majority of tissues (Supp. Fig. 8a), most significant associations were specific to only 1-3 tissues (Bonferroni-corrected p < 0.05; Fig. 4a, Supp. Fig. 8b). Among genes positively associated with mutation load in multiple tissues (see Methods), we observed enrichments for GO categories that included nucleotide excision repair, cellular transport, cell adhesion, and macroautophagy; and categories including immune response, keratinization, and cell polarity for negative expression-mutation associations (Fig. 4b, Supp. Tables 8,9). Consistent with these results, several of these same processes (e.g. cellular transport, autophagy and cell adhesion) are known to be associated with mutation load in cancer or normal cells^32–34^.

**Figure 4.**
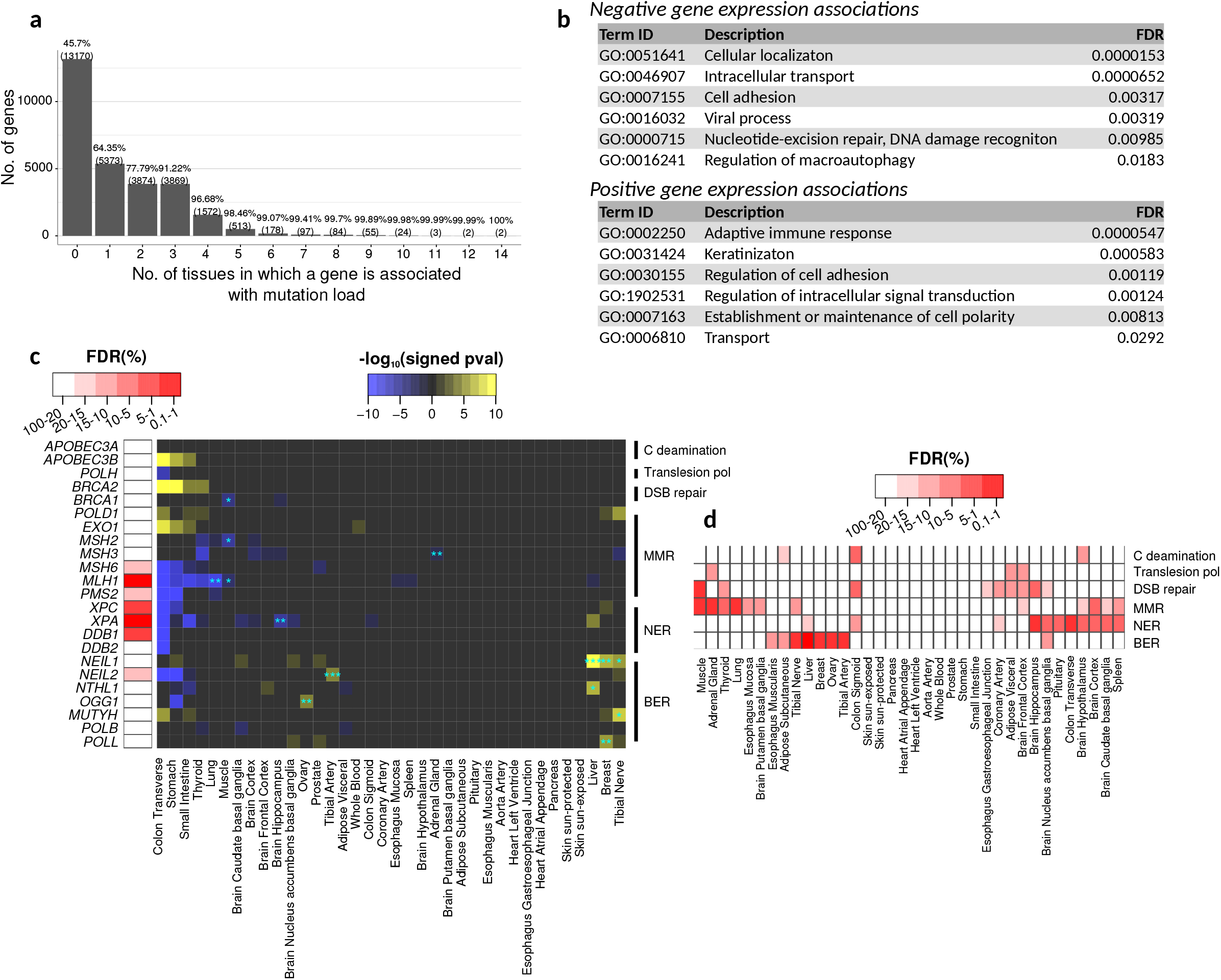
Mutation load is associated with the expression of genes and pathways. **a,** Histogram of the number of tissues for which each gene was significantly (Bonferroni corrected p < 0.05) associated with mutation load. Associations were estimated for each tissue using linear models controlling for population structure and biological and technical cofactors (see Methods). **b,** Genes whose expression was negatively (top) or positively (bottom) associated with C>T mutation load in multiple tissues are enriched in these representative GO categories (see Methods and Supp. Tables 8,9). **c,** Individual gene-tissue associations between C>T mutations and expression of genes involved in DNA repair or DNA mutagenesis (right panel). Blue asterisks denote significant associations using a permutation-based FDR strategy (see Methods; * FDR < 0.2, ** FDR < 0.1, *** FDR < 0.05). Shown on the left panel are genes whose expression was associated with mutation load across all tissues more than expected by chance at the indicated FDR (see Methods). **d,** Group-level gene expression associations of the shown pathways and C>T mutations across tissues (see Methods). Heatmap columns in c and d are ordered based on a hierarchical clustering.

To further explore the contribution of different DNA repair pathways to mutagenesis, we focused on the associations between the expression of repair pathway members with mutation load. Specifically, we analyzed genes involved in double-strand break (DSB) repair, mismatch repair (MMR), nucleotide-excision repair (NER), base excision repair (BER), the DNA deaminases *APOBEC3A/B* and the translesion polymerase *POLH*. We found 14 associations at a <20% FDR, 8 at <10% FDR, and 2 at <5% FDR (Fig. 4c blue asterisks; Supp. Fig. 8c-g) which are further described in Supplementary Note 2.

We then tested genes in these pathways for consistent associations with mutation load across tissues (see Methods; Fig. 4c, left panel; Supp. Fig. 10a-d). We observed significant hits from NER (*XPA*, FDR=0.01; *XPC*, FDR=0.07; *DDB1*, FDR=0.07), MMR (*MLH1*, FDR=0.05; *MSH6*, FDR=0.16; *PMS2* FDR=0.17), and BER (*NEIL2*, FDR=0.15). The associations were generally in the expected direction, based on what is known about each gene (see Supp. Note 3); for example, all associations of *MLH1* were negative, indicating lower expression associated with higher mutation load (Fig. 4c, Supp. Fig. 8e). *MLH1* silencing contributes to cancer development and mutagenesis^35^, and our results suggest that natural variation of *MLH1* expression is associated with mutagenesis across many tissues in the same direction as in cancer.

To further explore factors that may contribute to mutation load, we analyzed the expression of entire pathways or functionally related gene sets in each tissue (see Methods). MMR, BER, and NER are all strongly associated with mutation load in multiple tissues, but with very little overlap among them (Fig. 4d; Supp. Fig. 10e-i). As a result, these associations are primarily dominated by just one or two pathways per tissue.

In summary, most transcriptional signatures associated with mutation load are tissuespecific, however, genes associated with mutation load in several tissues are enriched in a number of pathways including DNA repair. These results paint a complex landscape where mutational load across non-disease tissues is associated with a variety of cellular functions.

### Cancer driver genes are enriched for mutations and under positive selection in non-disease tissues

To investigate the extent to which cancer-associated mutations exist in a pre-cancerous state, we calculated the enrichment of all COSMIC^36^ cancer point mutations in our mutation maps (see Methods). Many samples were highly enriched (Hypergeometric Bonferroni-corrected p<0.05), with some having more than 20% overlap with COSMIC mutations (Fig. 5a). In contrast, we observed almost no significant overlaps in a negative control (using a permuted set of mutations per-sample, that conserves mutation frequencies and genomic regions from the original set of mutations; see methods; Supp. Fig. 11a). Sun-exposed skin had the highest level of overlap (Hypergeometric BH-FDR<0.05), with 100% of samples having significant enrichments, followed by sun-protected skin and skeletal muscle (Fig. 5a). We observed the lowest overlaps across the seven brain regions, aorta, and spleen.

**Figure 5.**
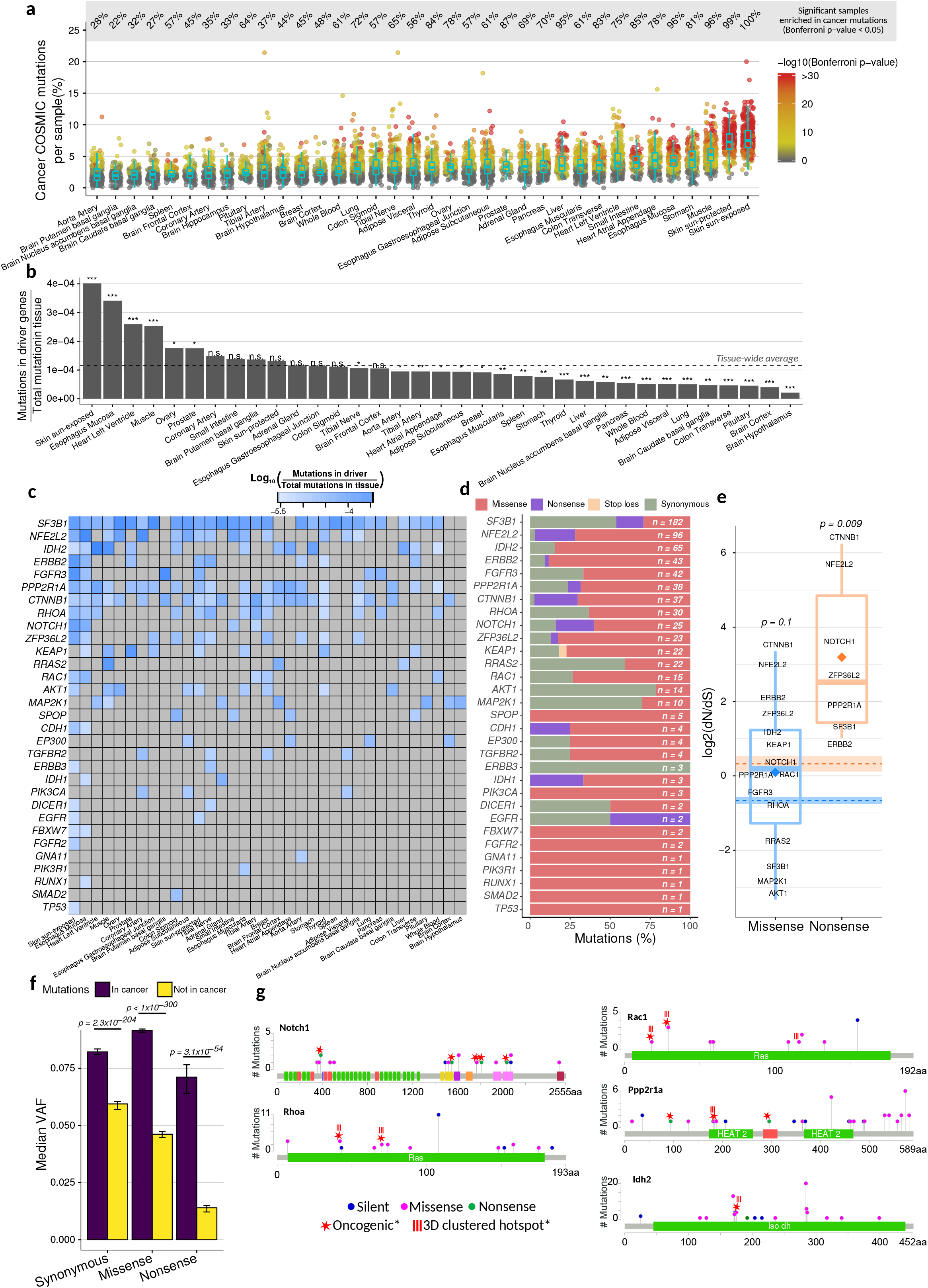
Cancer driver genes evolve under strong positive selection, and cancer mutations are enriched in healthy tissues. **a,** Percentage of COSMIC cancer mutations observed per sample and grouped by tissue; p-values for enrichment were calculated using a hypergeometric test accounting for sequencing coverage, total number of mutations per sample, and total number of COSMIC mutations, and the three possible alternate alleles that any given reference allele can have (see Methods). P-values are Bonferroni-corrected across all samples. **b,** Relative mutation rates of a selected group of 53 genes known to carry cancer driver mutations^37^ (only 31 of them had at least 1 mutation in this study); the tissue-wide average is indicated with the dotted line. Significant deviation from the tissue-wide average was calculated using the binomial distribution and the tissue-wide average mutation rate. Benjamini-Hochberg FDR: *** (FDR < 0.001), ** (FDR < 0.01), * (FDR < 0.05). **c,** Individual mutation rates for each cancer driver gene across all tissues. **d,** Percentage of mutations for each cancer driver gene stratified by impact to amino acid sequence; n is the total number of mutations observed in a gene. **e,** dN/dS values for missense (blue) and nonsense mutations (orange) in cancer driver genes calculated using dndsloc^9,38^ (see Methods); averages per group are shown as rhomboids and their respective genome-wide averages are shown as dashed lines along with their 95% confidence intervals after bootstrapping 10,000 times. P-values indicate the probability of observing a higher average dN/dS from 10,000 equally-sized randomly-sampled groups of genes (see Methods). **f,** Median variant allele frequency (VAF) for each mutation type based on their impact to the amino acid sequence and colored by their cancer status. Mutations “in cancer” (purple bars) are those that overlap with the COSMIC database in both base change and position, and mutations “not in cancer” (yellow bars) are those that overlap with COSMIC only in position but not in base change. Error bars represent the 95% confidence interval after bootstrapping 1000 times; p-values are from two-sided Mann-Whitney tests. **g,** Mutation maps of five cancer driver genes; oncogenic state was obtained from oncoKB^39^, and clustered mutation annotations were obtained from the databases Cancer Hotspots^42^ and 3D Hotspots of mutations occurring in close proximity at the protein level^43^.

Focusing our analysis on a panel of 53 known cancer driver genes^37^ (Supp. Table 10), we observed mutations in 31 of them (after excluding potential false-positive mutations; see Methods and Supp. Table 11). Some tissues showed significantly greater or lower mutation rates across these genes (Fig. 5b). Sun-exposed skin was once again the most significant tissue, with several others at FDR < 0.05, such as muscle and heart (Fig. 5b). Interestingly, *MAP2K1* and *RRAS2* are highly mutated in muscle, and *IDH2* and *PPP2R1A* are highly mutated in both heart and skeletal muscle (Fig. 5c). Conversely, several tissues—including most brain regions—had significantly lower mutation rates than average in these cancer drivers (Fig. 5b).

Observed mutations in cancer driver genes were then classified by their impact on protein sequences. We found that some genes were dominated by missense mutations, such as *IDH2*, while others showed an excess of synonymous mutations, such as *MAP2K1* (Fig. 5d).

To assess the selective pressures acting on these genes we used a dN/dS approach, which assesses the ratio of non-synonymous to synonymous mutations while accounting for sequence composition and variable mutation rates across the genome^38^ (see Methods). dN/dS values close to 1 represent little or no detectable selection, dN/dS>1 suggests positive selection, and dN/dS<1 suggests purifying selection. Our analysis showed that most of these cancer drivers are evolving under positive selection (Fig. 5e). Missense mutations had a mean dN/dS=1.06 whereas nonsense mutations had a mean dN/dS=9.14 (indicating over 9-fold enrichment for nonsense mutations compared to the expectation based on synonymous mutations in the same genes); the latter is significantly more than expected from genome-wide values (permutation-based p=0.1 and p=0.009, respectively; Fig. 5e).

*NOTCH1* has been recently observed to evolve under positive selection in non-cancerous skin^15^ and esophagus^18,19^. Consistent with this, we found the highest rates of *NOTCH1* mutations in these same two tissues (Fig. 5c), with strong signals of positive selection (*NOTCH1* missense dN/dS = 1.46, nonsense dN/dS = 14.04). We also observed *NOTCH1* mutations at slightly lower rates in the small intestine and tibial artery (Fig. 5c).

To further explore the effects of selection on cancer-associated mutations, we performed a VAF analysis (similar to Fig. 2d). Using the large catalog of COSMIC cancer point mutations (as in Fig. 5a), we created two subsets of our somatic mutation list: those that have been observed in cancer samples, and those at precisely the same location as a known cancer mutation, but with a different base change (which act as well-matched negative controls). We then separated each of these two subsets into three mutation types: synonymous, missense, and nonsense. In all three cases, the VAFs of the cancer-associated mutations were significantly higher than the matched non-cancer-associated ones (Fig. 5f). Using the synonymous non-cancer mutations as representative of neutrality (similar to dN/dS), we found that all three cancer-associated mutation types had higher VAF than these neutral proxies (Fig. 5f), suggesting the action of positive (and not just weaker negative) selection. Interestingly, the ratio of cancer/non-cancer mutation VAFs was lowest for synonymous (1.4-fold), intermediate for missense (2.0-fold), and highest for nonsense (5.2-fold), suggesting stronger positive selection for more extreme mutations. These results are consistent with the dN/dS analysis in Fig. 5e, which also showed the strongest positive selection on nonsense mutations in cancer driver genes, but the VAF analysis has substantially more power due to the greater number of mutations analyzed.

We also found a number of specific mutations manually curated as oncogenic or likely oncogenic from OncoKB^39^ (Supp. Table 12). For instance, all nonsense mutations that we observed in *NOTCH1* have been labelled as oncogenic (Fig. 5g). We found many other oncogenic mutations as well; for example, *RHOA* and *RAC1* had oncogenic mutations in their Ras domains (Fig. 5g).

Together these results show that cancer mutations are enriched in non-disease tissues and not only do they accumulate in driver genes, but the majority of these genes are evolving under positive selection, suggesting that these mutations increase cellular proliferation well before any cancer is observed.

## DISCUSSION

We developed a method to detect rare somatic mutations from RNA-seq data and applied it to over 7,500 tissue samples (Fig. 1). To our knowledge this is the largest map to date of somatic mutations in non-cancerous tissues.

It has been proposed that somatic mutations contribute to aging and organ deterioration^40^; consistent with this, we observed a positive correlation between age and mutation burden in most tissues. Interestingly, several brain regions are among the tissues exhibiting stronger age correlation, and somatic mutations have been shown to have a role in neurodegeneration^24^.

We observed largely tissue-specific behaviors and some pervasive observations shared across tissues. Mutation profiles are defined by their tissue-of-origin, mutation maps are delineated by tissue-specific chromatin organization, and transcriptional signatures associated with mutation load are highly tissue-specific. These results suggest that different cell types are subjected to different evolutionary paths that could be dependent on environmental or developmental differences. For example, while most samples exhibit tissue-specific mutation profiles, some others like transverse colon and the small intestine have similar profiles. Additionally, we observed that genes whose expression is associated with mutation load in several tissues are enriched in DNA repair, autophagy, immune response, cellular transport, cell adhesion and viral processes; and while these functions have been implicated in mutagenesis in cancer^32–34^, our results highlight how expression variation of these genes associates with mutational variation in healthy tissues.

Cancer mutations are enriched across many organs. Muscle and heart tissue are particularly interesting because they have lower-than-expected mutation rates, but those mutations are highly enriched for cancer mutations and had high mutation rates in cancer drivers. Sarcomas are tumors originated from soft tissues – including muscle – that are relatively uncommon and have a low density of point mutations but high copy number variation^41^. Our results show that low mutation rates are also observed in these soft tissues, but compared to tumors, cancer mutations are frequent. The functional implications of this observation will need to be explored in further detail.

Positive selection of driver genes has been recently observed in healthy tissues^15,18^. Accordingly, we confirmed that *NOTCH1* is under positive selection in skin and esophagus. We found other genes positively selected both broadly (e.g. *IDH2, CTNNB1, NFE2L2*) and with more tissue-specific patterns (e.g. *KEAP1* in prostate, thyroid and muscle; *RAC1* in skin, esophagus, and breast), which will be important subjects for future studies.

In general, mutations previously observed in cancer studies were found in high abundance across many healthy tissues, and our VAF analysis showed signatures of positive selection acting on them. In contrast, when looking at all mutations and in particular those not previously seen in cancer, we found that missense and nonsense mutations are generally under negative selection. Our results reconcile recent studies reporting prevalent positive^9^ or negative^11^ selection in somatic evolution, as we find evidence of both co-existing in different sets of mutations.

Our findings paint a complex landscape of somatic mutation across the human body, highlighting their tissue-specific distributions and functional associations. The prevalence of cancer mutations and positive selection of cancer driver genes in non-diseased tissues suggests the possibility of a poised pre-cancerous state, which could also contribute to aging. Finally, our method for inferring somatic mutations from RNA-seq data may help accelerate the study of somatic evolution and its role in aging and disease.

## SUPPLEMENTARY TABLES

**Supplementary Table 1.** Average percentage elimination of putative mutation calls by persample false-positive filters across all tissues.

**Supplementary Table 2.** Total number of samples per tissue included for the final set of mutation calls.

**Supplementary Table 3.** List of all somatic mutations identified in this study.

**Supplementary Table 4.** P-values (−log_10_[p-value]) for the coefficients of each feature used in a linear regression on the total number of mutations per tissue.

**Supplementary Table 5.** Average percentage of each mutation type across samples of the given tissue.

**Supplementary Table 6.** Significant associations between biological sex and mutation load across tissues and mutation types.

**Supplementary Table 7.** Tissue correspondence between the GTEx^23^ and Roadmap Epigenomics projects^28^.

**Supplementary Table 8.** Significant GO enrichments for genes whose expression is significantly and negatively associated with C>T mutation load across several tissues.

**Supplementary Table 9.** Significant GO enrichments for genes whose expression is significantly and positively associated with C>T mutation load across several tissues.

**Supplementary Table 10.** List of cancer driver genes used in this study^37^.

**Supplementary Table 11.** List of mutations in cancer driver genes after further elimination of potential false positives.

**Supplementary Table 12.** List of mutations in cancer driver genes annotated in Oncokb^39^.

**Supplementary Table 13.** Correspondence between GTEx^23^ tissues and cancer types from Tomasetti and Vogelstein^44^.

**Supplementary Figure 1.**
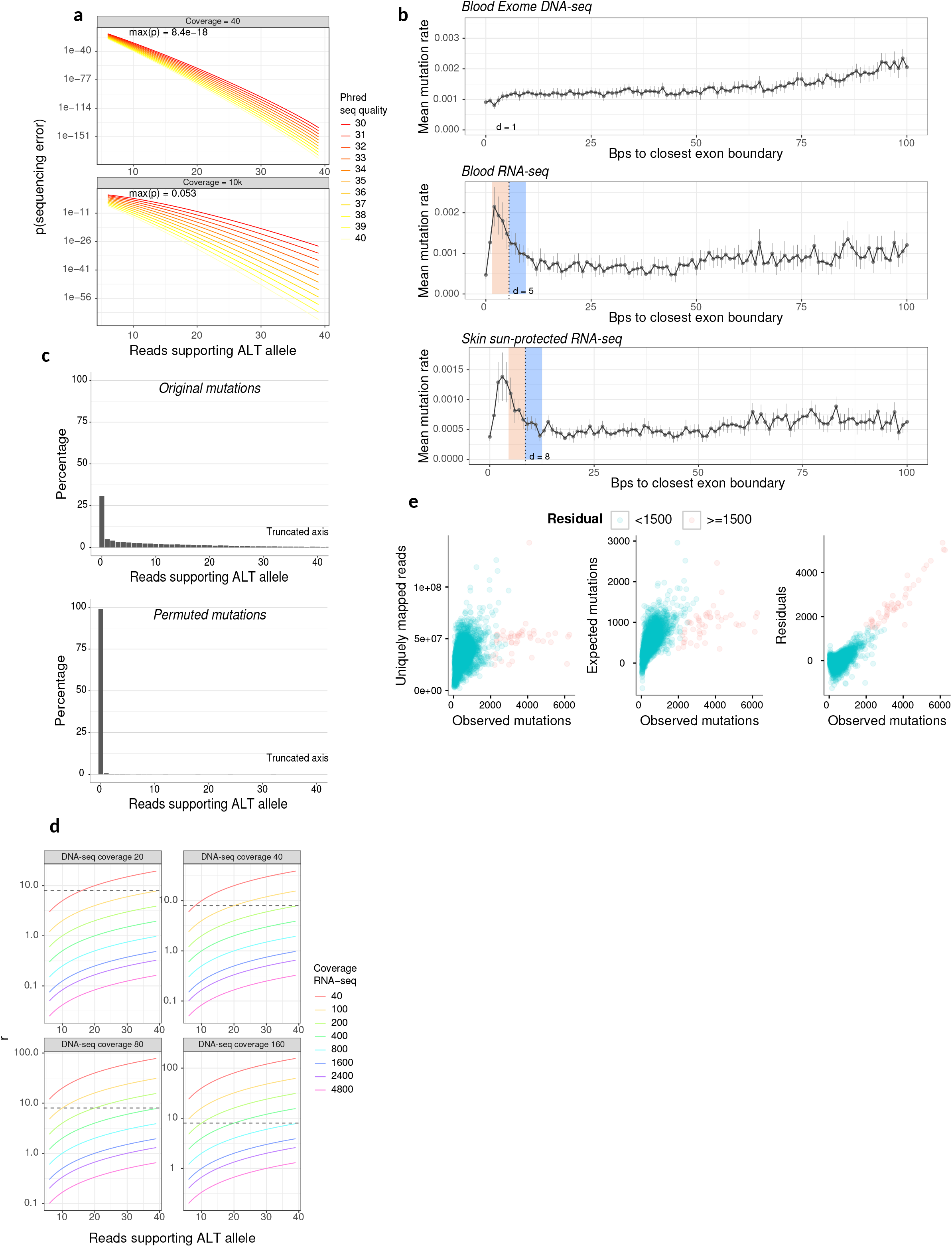
Statistics associated to a method for calling DNA mutations from RNA-seq data. **a,** Probabilities of observing a sequencing error (y axis) on a position covered with the lowest coverage used in this study (top panel, 40 reads), or a highly covered base (bottom panel, 104 reads), with increasing number of reads supporting the alternate allele (x axis), at different phred sequencing scores (colored lines). Probability is calculated using the right tail of a binomial distribution on the number of observed reads supporting a mutation [X ~ binom(n = coverage, p=10^−Prehd/10^x1/3); Phred-based sequencing error probability is multiplied by 1/3 to account for three potential different bases to mutate]. **b,** Average mutation rate is high close to splice junctions in RNA-seq (bottom 2 rows) but not in DNA-seq data (top row). Mutation rate was calculated by taking the number of mutations observed at a given distance from an exon junction and dividing it by the number of reads covering positions at that distance. Vertical dashed lines represent the point of inflection at which mutation rate stabilizes; we used a 1-bp sliding window to identify the position with maximum absolute difference between mutation rates of the four bps downstream and the four bps upstream of that position. Error bars are the 95% confidence intervals based on bootstrapping 1,000 times. **c,** DNA somatic mutations called from RNA-seq were validated by assessing the percentage of mutations for which there was at least one read supporting the alternate allele in matched exome DNA-seq data (see Methods). The top panel shows the histogram of number of reads supporting the alternate allele in DNA-seq for all mutations found in RNA-seq. The bottom panel shows the same data after randomly assigning an alternate allele to the mutations called in RNA-seq, effectively creating a distribution expected by chance (see Methods). X axes were truncated for visual purposes. **d,** For the method validation, we compared RNA-seq-based mutation calls to DNA-seq data. To address differences in coverage between the two methods we only compared positions with r >= 8 (dashed line; r effectively represents the number of expected reads that support the alternate allele in DNA-seq at a position given the alternate allele frequency observed in RNA-seq at that position and the DNA-seq coverage; see Methods), which ensures considering only positions for which reads supporting the alternate allele were expected to be found in DNA-seq given the coverage of that position in both experiments. **e,** Hyper-mutated samples were identified by applying a linear regression between the number of mutations and sequencing depth (Uniquely mapped reads), age, BMI and gender. The observed number of mutations was mostly explained by sequencing depth (left panel) and the expected number of mutations agrees for most samples with the observed number mutations (middle panel). Hyper-mutated samples were tagged as the ones with residuals from the linear regression of >=1,500, meaning they had at least 1,500 mutations more than expected (right panel; see Methods for more details).

**Supplementary Figure 2.**
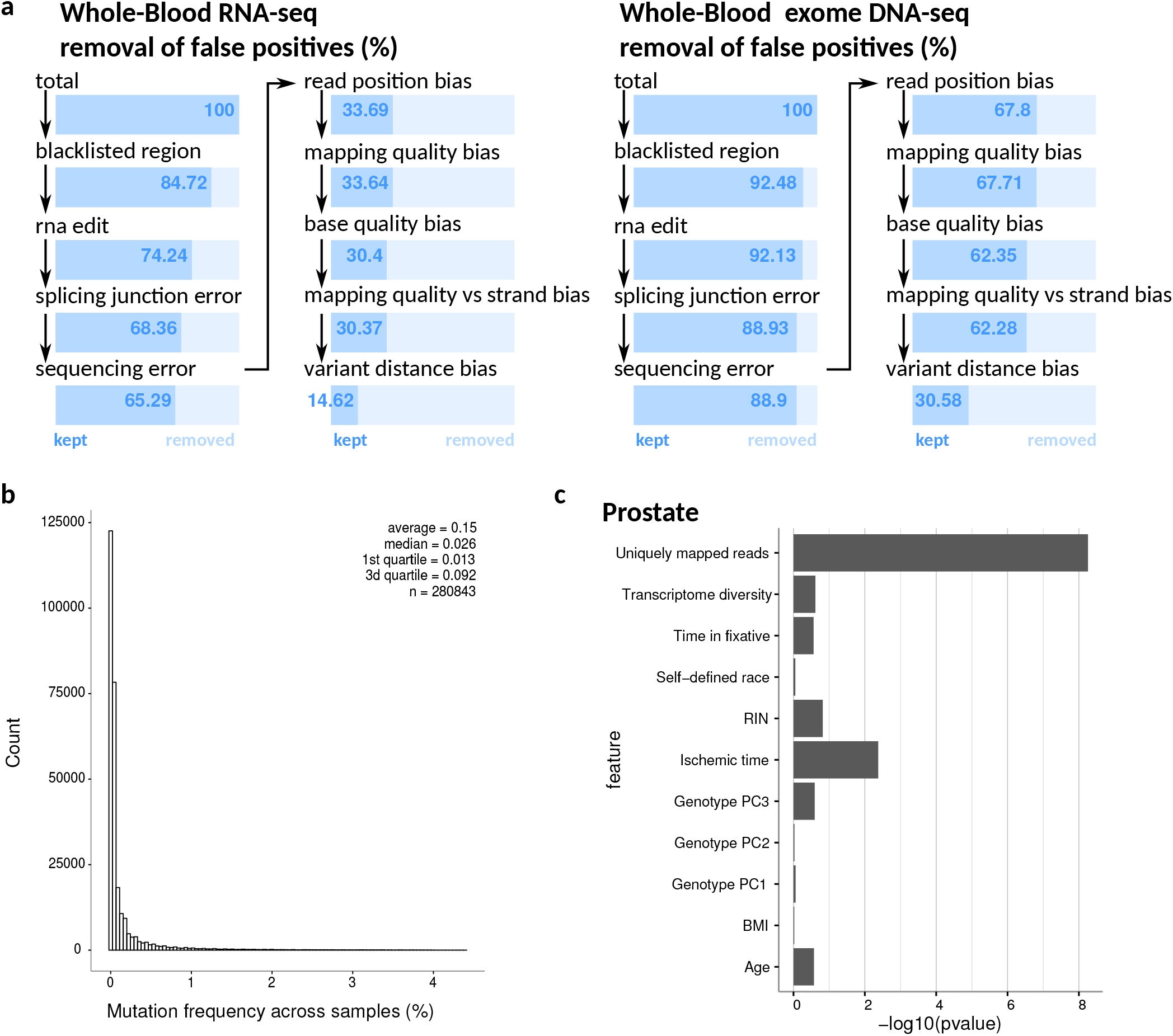
Calling of somatic DNA-mutations in the GTEx cohort. **a,** Per-sample filters applied to mutation calls in blood RNA-seq (left) and DNA-seq (right) (See Supp. Table 1 for all-tissue data). **b,** Frequency histogram of unique mutations across samples. **c,** Significance of associations between total number of mutations observed in prostate with biological and technical variables. P-values were obtained from a linear regression (see Methods; see Supp. Table 4 for all-tissue data).

**Supplementary Figure 3.**
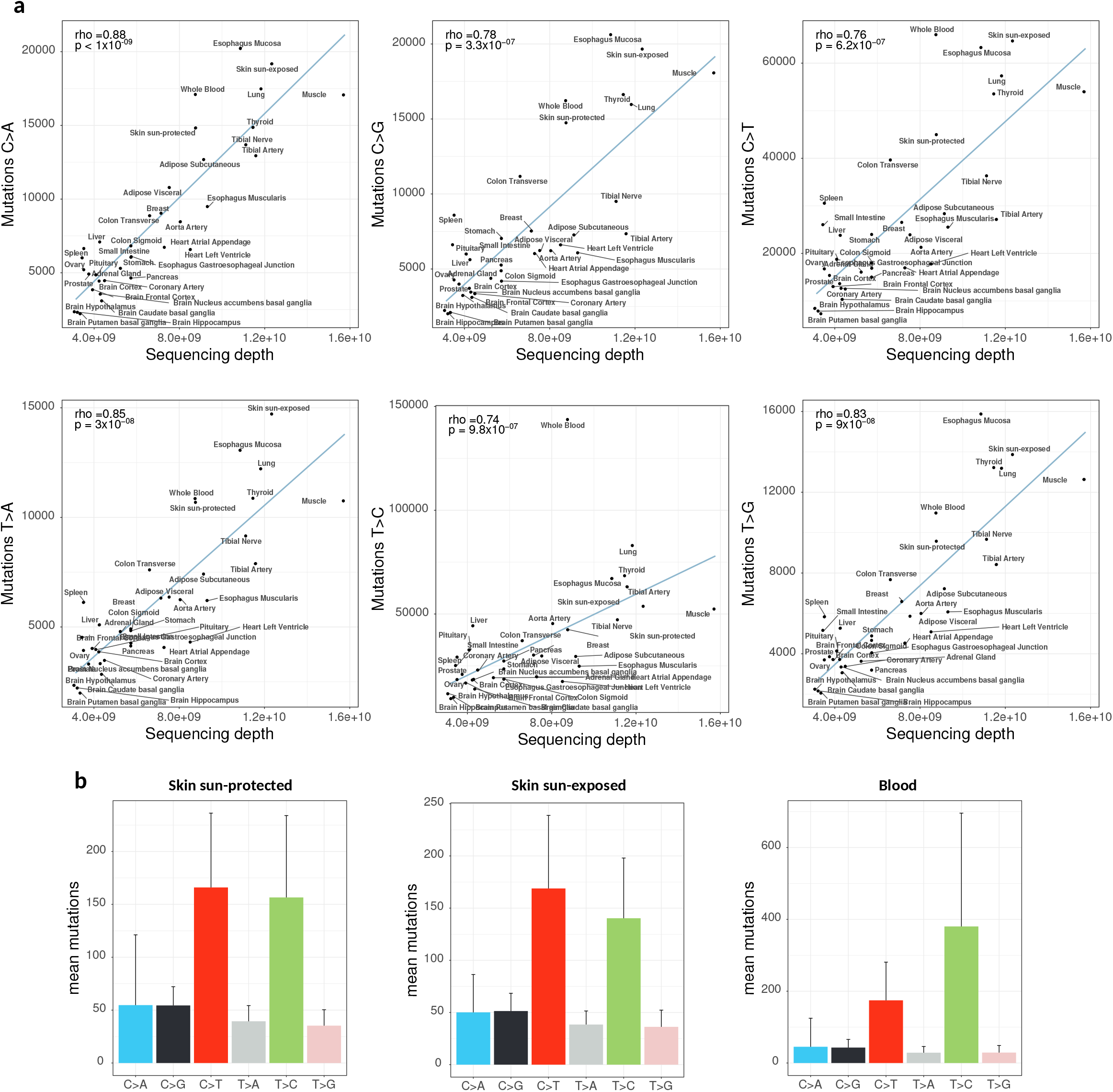
Mutation load across different mutation types in non-disease human tissues. **a,** Across all mutation types, the total number of mutations observed in a tissue is explained by the total sequencing depth of that tissue. A linear regression line is shown in blue; tissues above it exhibit more mutations than expected by sequencing depth and tissues below it show fewer mutations than expected. Rho is the Spearman coefficient. **b,** Representative examples for relative contributions of different mutation types across different tissues (see Supp. Table 5 for all-tissue data).

**Supplementary Figure 4.**
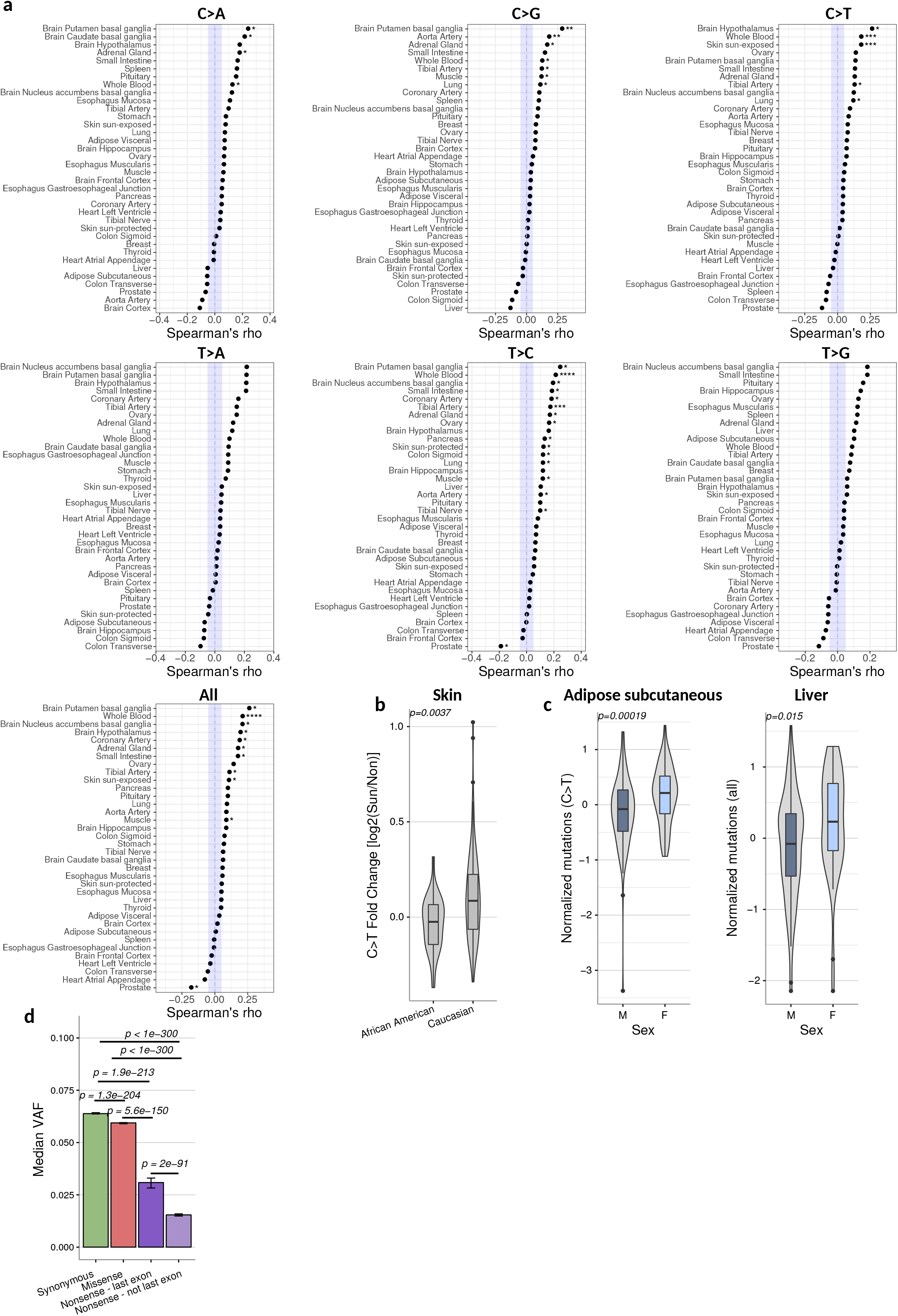
Phenotypic associations and properties of mutation load in the human body. **a,** Age associations (x axis) with mutation load per tissue (y axis) across mutation types (panels). P-values were obtained by assessing the fraction of permutation-based rho values greater than the original rho value for positive rho values, or the fraction of permutation-based rho values smaller than the original rho value for negative rho values. A total 10,000 permutations were performed. FDRs were obtained for each panel using the Benjamini-Hochberg method: **** (FDR < 0.01), *** (FDR < 0.05), ** (FDR < 0.1), * (FDR < 0.2). **b,** Fold-change of C>T mutations between matched sun-exposed and sun-protected skin samples from the same individual and stratified by self-reported race. P-value is based on a two-sided Mann-Whitney test. **c,** Two representative examples of associations between mutation load and biological sex. To control for sequencing depth and other technical artifacts, mutation values were obtained as the residuals from a linear regression using technical features as explanatory variables (see Methods, see Supp. Table 6 for all significant associations with sex). **d,** Related to Figure 2d, median variant allele frequency (VAF) across all mutations for each mutation type based on their impact to the amino acid sequence; nonsense mutations were divided into two different groups based on whether they are located in the last exon; error bars represent the 95% confidence interval after bootstrapping 1000 times; p-values are from two-sided Mann-Whitney tests.

**Supplementary Figure 5.**
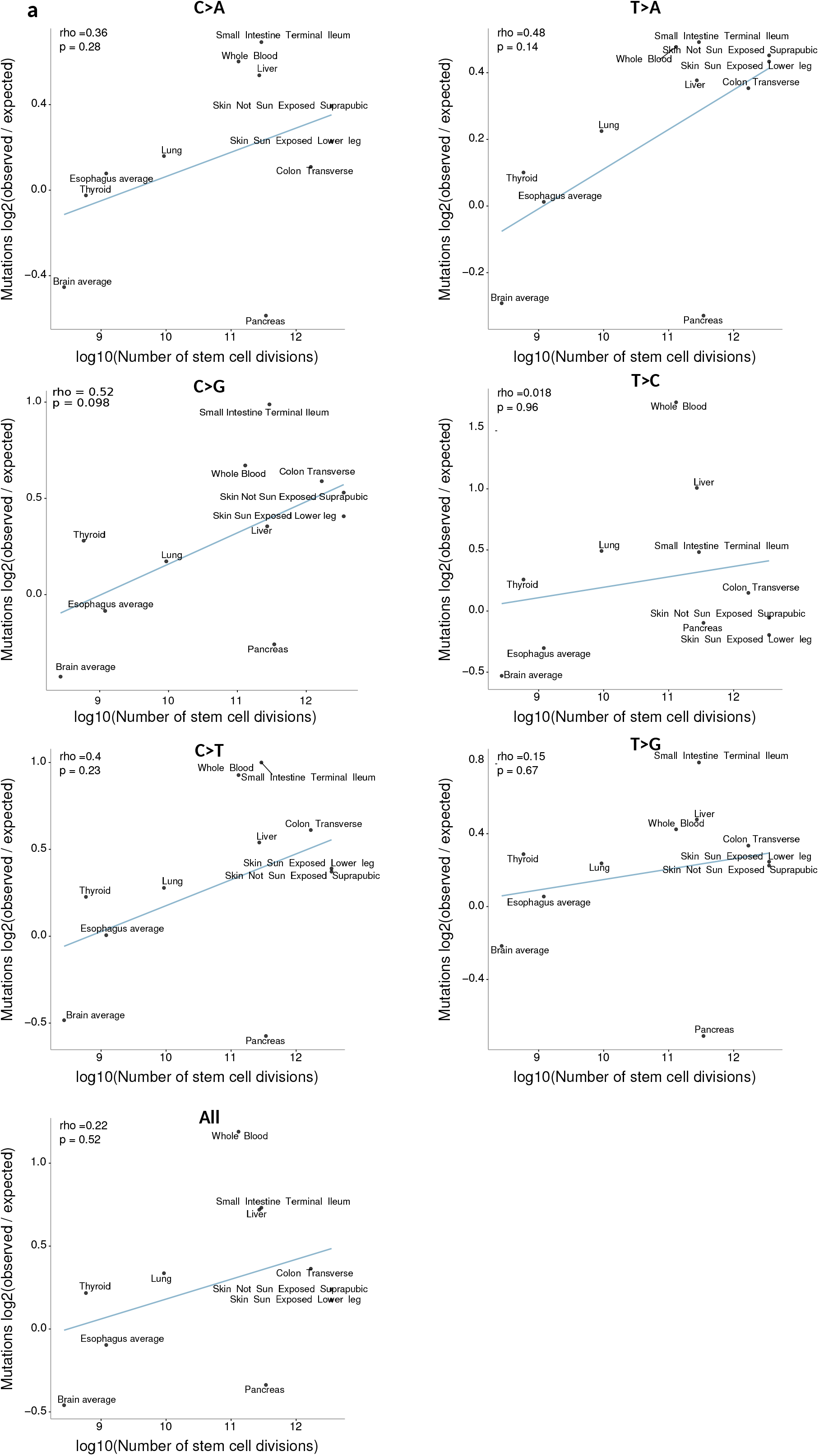
Number of stem cell divisions correlates weakly with mutation load in human tissues. Supplementary Figure 5. Number of stem cell divisions correlates weakly with mutation load in human tissues. Associations between different mutation types and the number of stem cell divisions of the most abundant cell type for each tissue^44^. Observed/expected mutations (y-axis) were calculated by dividing the number of mutations observed in a given tissue by the predicted number of mutations based on the sequencing depth of the tissue from the linear regression in Figure 2a. Associations between different mutation types and the number of stem cell divisions of the most abundant cell type for each tissue^44^. Observed/expected mutations (y-axis) were calculated by dividing the number of mutations observed in a given tissue by the predicted number of mutations based on the sequencing depth of the tissue from the linear regression in Figure 2a.

**Supplementary Figure 6.**
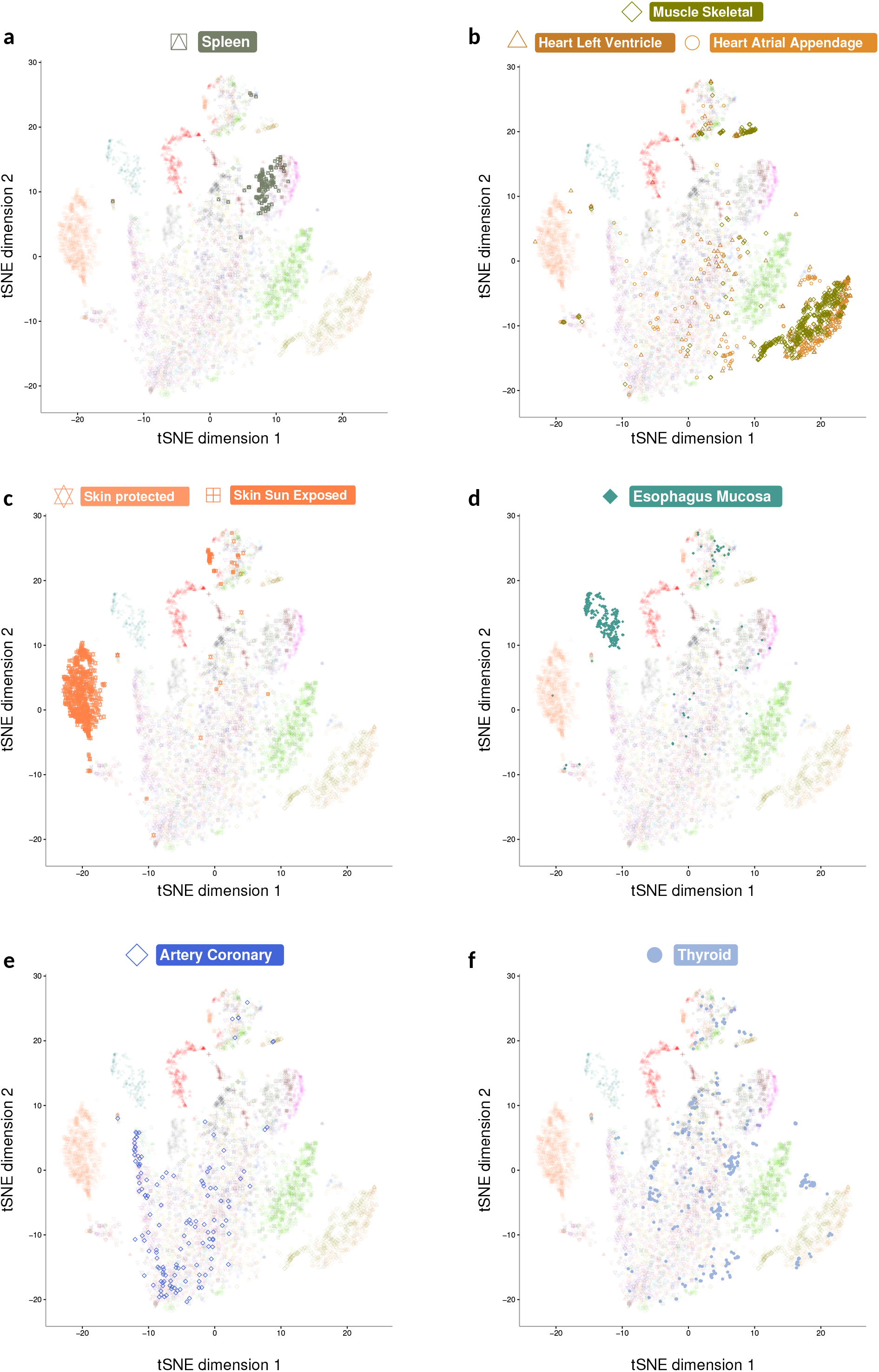
Mutation profiles cluster by tissue. **a-f,** tSNE plots as described in Figure 2e, highlighting individual (a,d) and grouped (b,c) tissues exhibiting highly similar within-tissue mutation profiles, and two tissues showing tissue profiles with weak clustering (e,f).

**Supplementary Figure 7.**
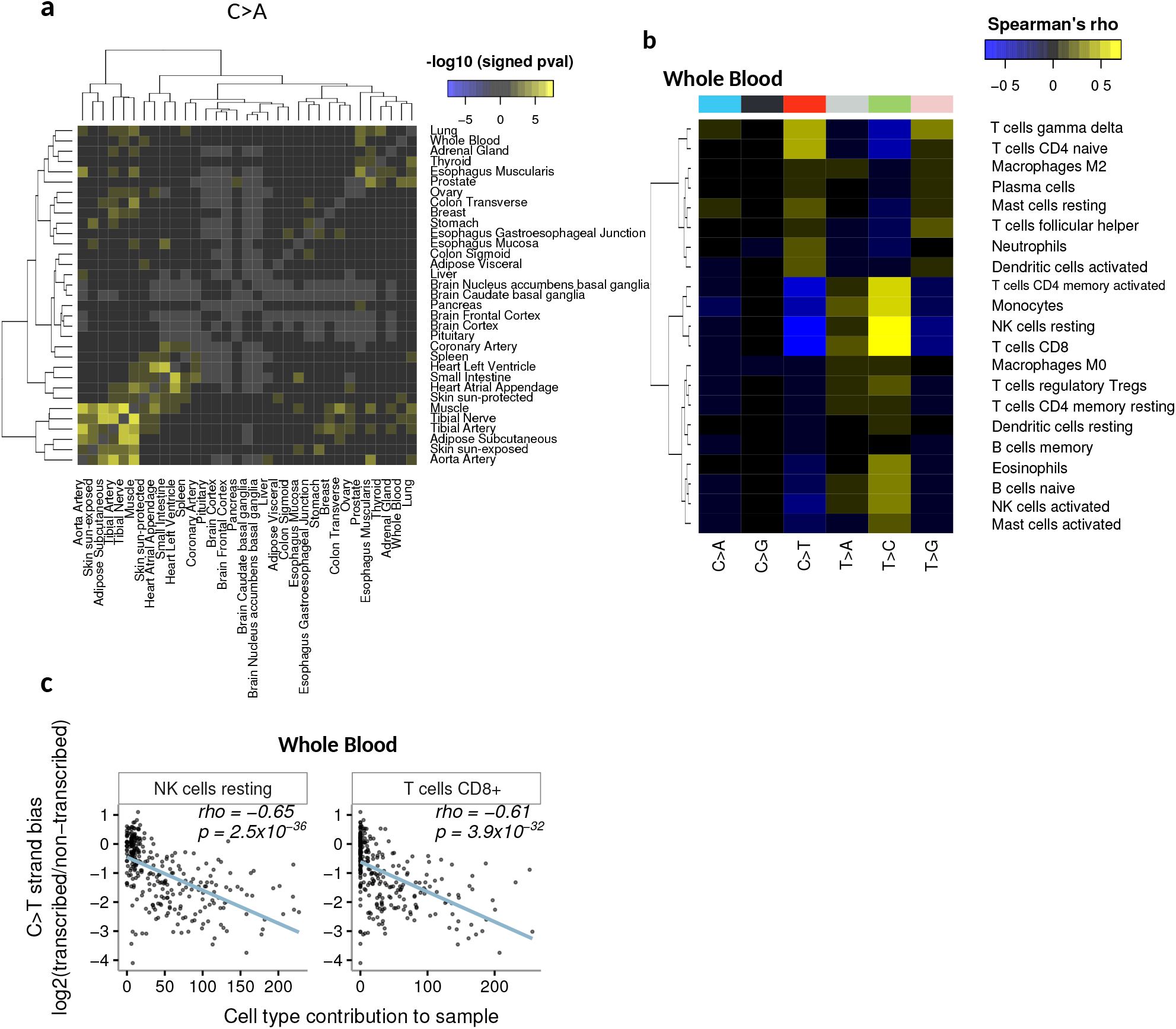
Inter-tissue mutational strand asymmetry correlations and cell type associations. **a,** C>A inter-tìssue strand asymmetry correlations; p-values are from Spearman correlations between strand asymmetry (ratio of mutations on the transcribed over the non-transcribed strand) in pairs of tissues from the same individual. Yellow indicates positive correlations, whereas blue indicates negative correlations. Tissues in light gray did not have enough samples to assess associations. Columns and rows were ordered using hierarchical clustering. **b,** Associations between C>T strand asymmetry (transcribed/ non-transcribed strand) and blood cell type composition. Cell type composition was obtained using CIBERSORT45 (see Methods). **c,** Top positive and negative correlations in blood (from panel b) between C>T strand asymmetry and cell type content.

**Supplementary Figure 8.**
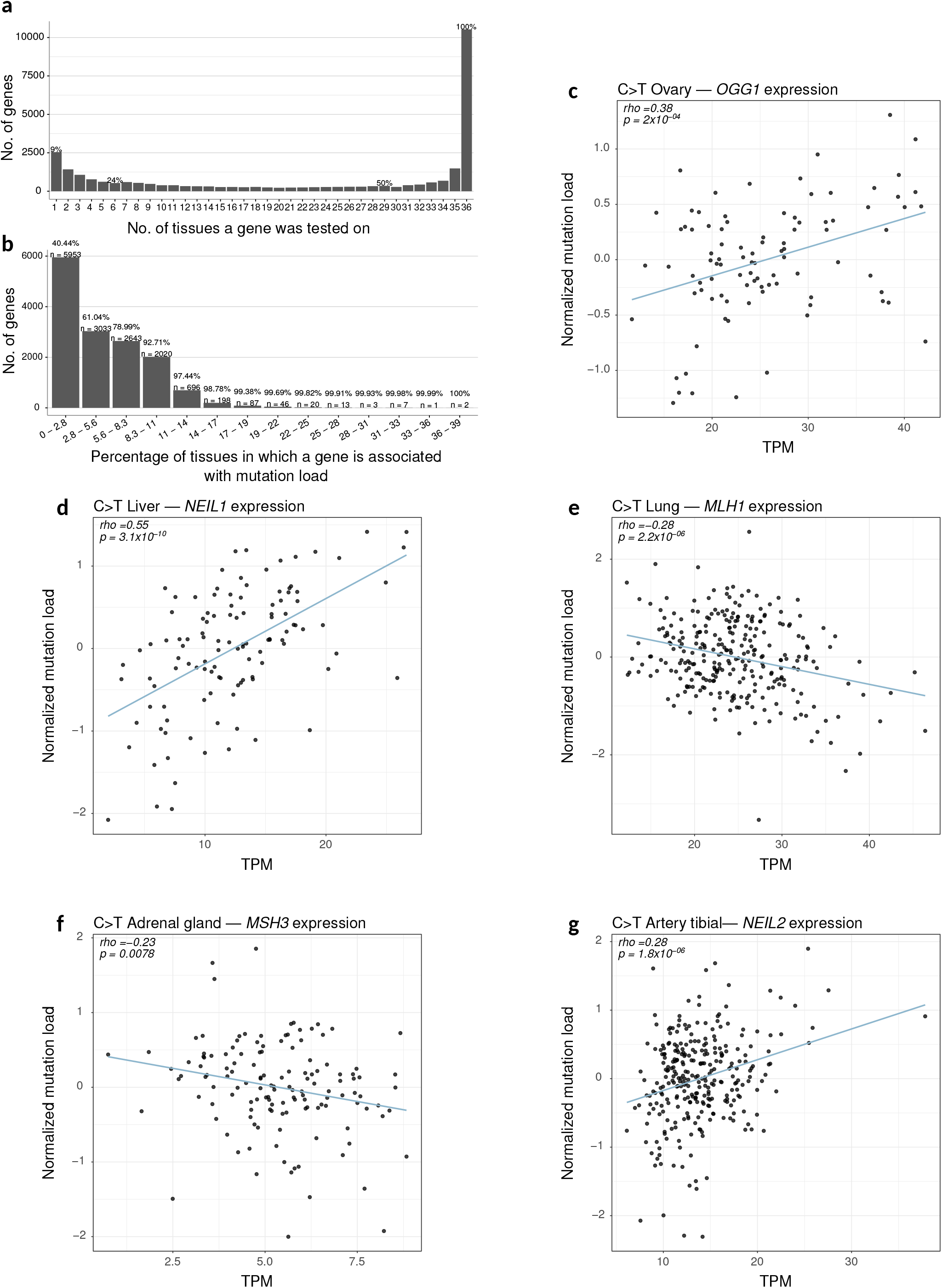
Gene expression associations with C>T mutation load. **a,** Histogram of the number of tissues in which a gene was tested for association between its expression and C>T mutation load. A gene was selected to be tested in a tissue based on having detectable expression in that tissue (see Methods). **b,** Histogram of the percentage of tissues exhibiting significant associations (p < 0.05 after Bonferroni correction) between expression of a gene and C>T mutation load. **c-g,** Examples of individual significant associations between C>T mutation load and expression of DNA repair genes in different tissues (0.001 < FDR < 0.2, see Fig. 4c). Mutation load is normalized by controlling for biological and technical factors (see Methods). Rho is the Spearman correlation coefficient.

**Supplementary Figure 9.**
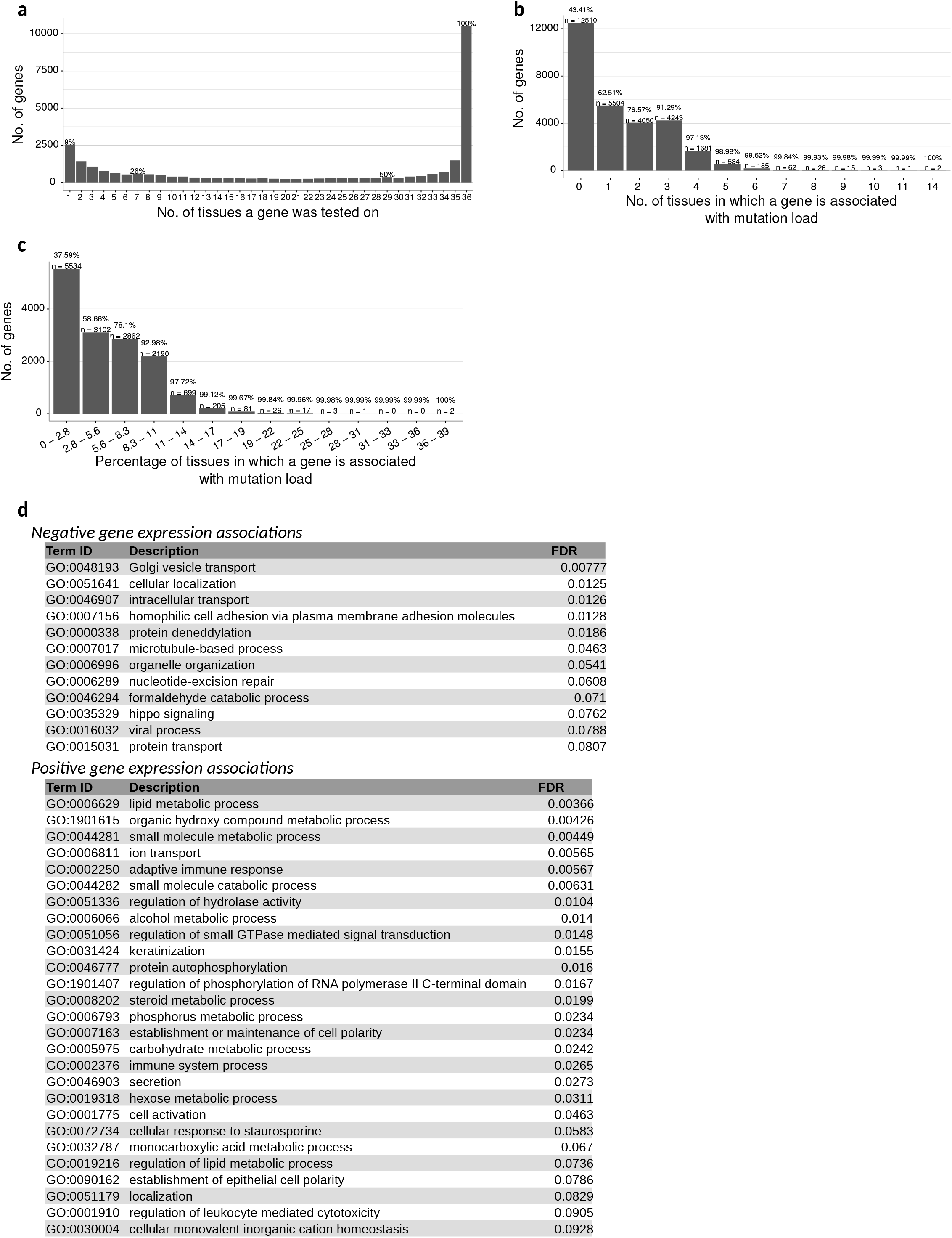
Gene expression associations with overall mutation load. Similar to Figure 4a,b but for all mutation types. **a,** Histogram of the number of tissues in which each gene was detectably expressed, and thus tested for association between its expression and mutation load across all mutation types (see Methods). **b-c,** Histogram of the number (b) or percentage (c) of tissues in which a gene was significantly associated with mutation load (p < 0.05, Bonferroni corrected). **d,** Genes whose expression is negatively (top) or positively (bottom) associated with mutation load in multiple tissues are enriched in these representative lists of GO categories (see Methods).

**Supplementary Figure 10.**
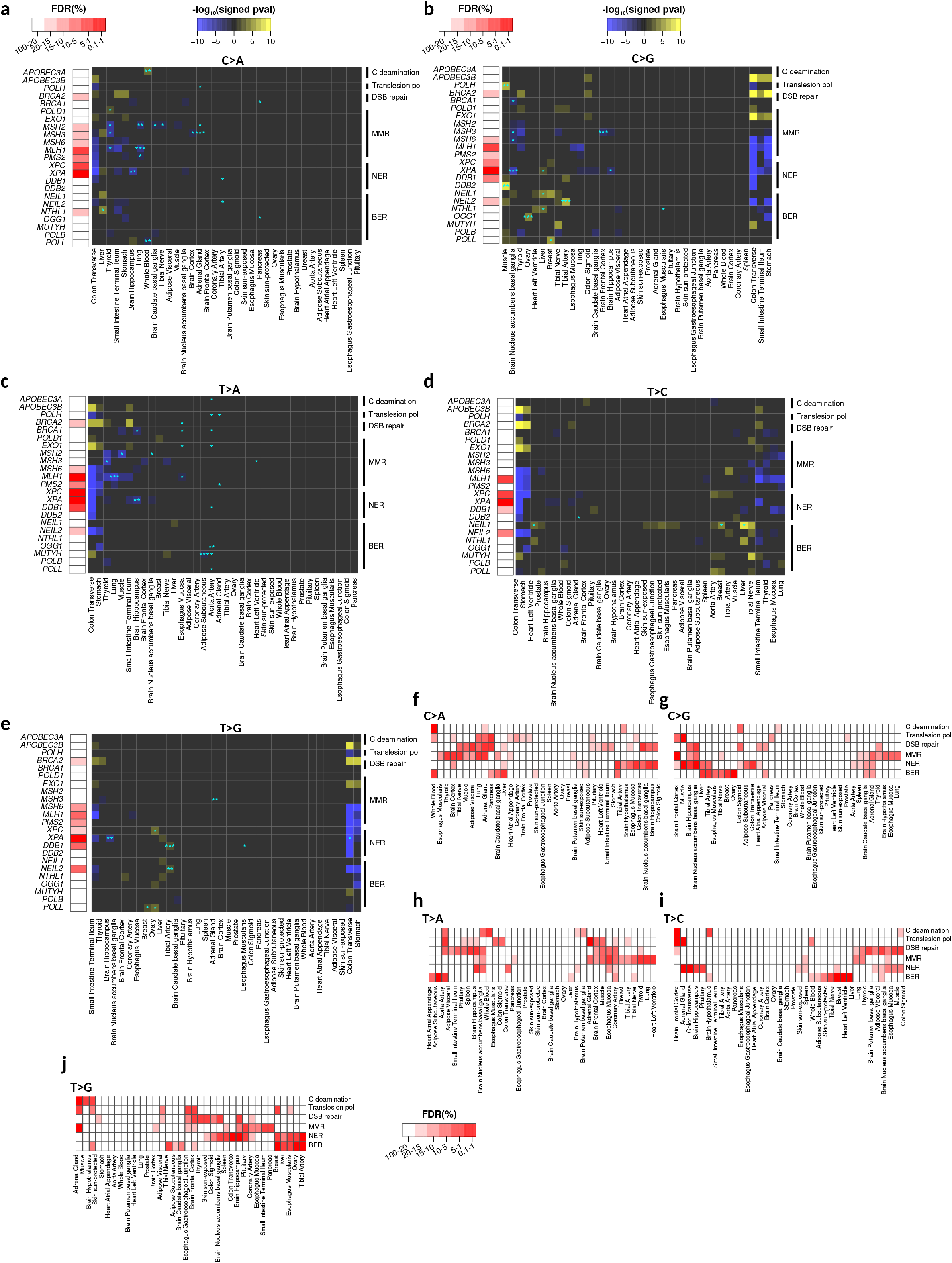
Mutation load associations with expression of genes involved in DNA repair or DNA mutagenesis. **a-d,** Individual gene-tissue associations between different mutation types and expression of genes involved in DNA repair or DNA mutagenesis (right panels); blue stars denote significant associations using a permutation-based FDR strategy (see Methods; * FDR < 0.2, ** FDR < 0.1, *** FDR < 0.05). Shown in the left panels are genes whose expression was associated across all tissues more than expected by chance at the indicated FDR (see Methods). **e-i,** Group-level gene expression associations of the shown pathways and different mutation types across tissues (see Methods).

**Supplementary Figure 11.**
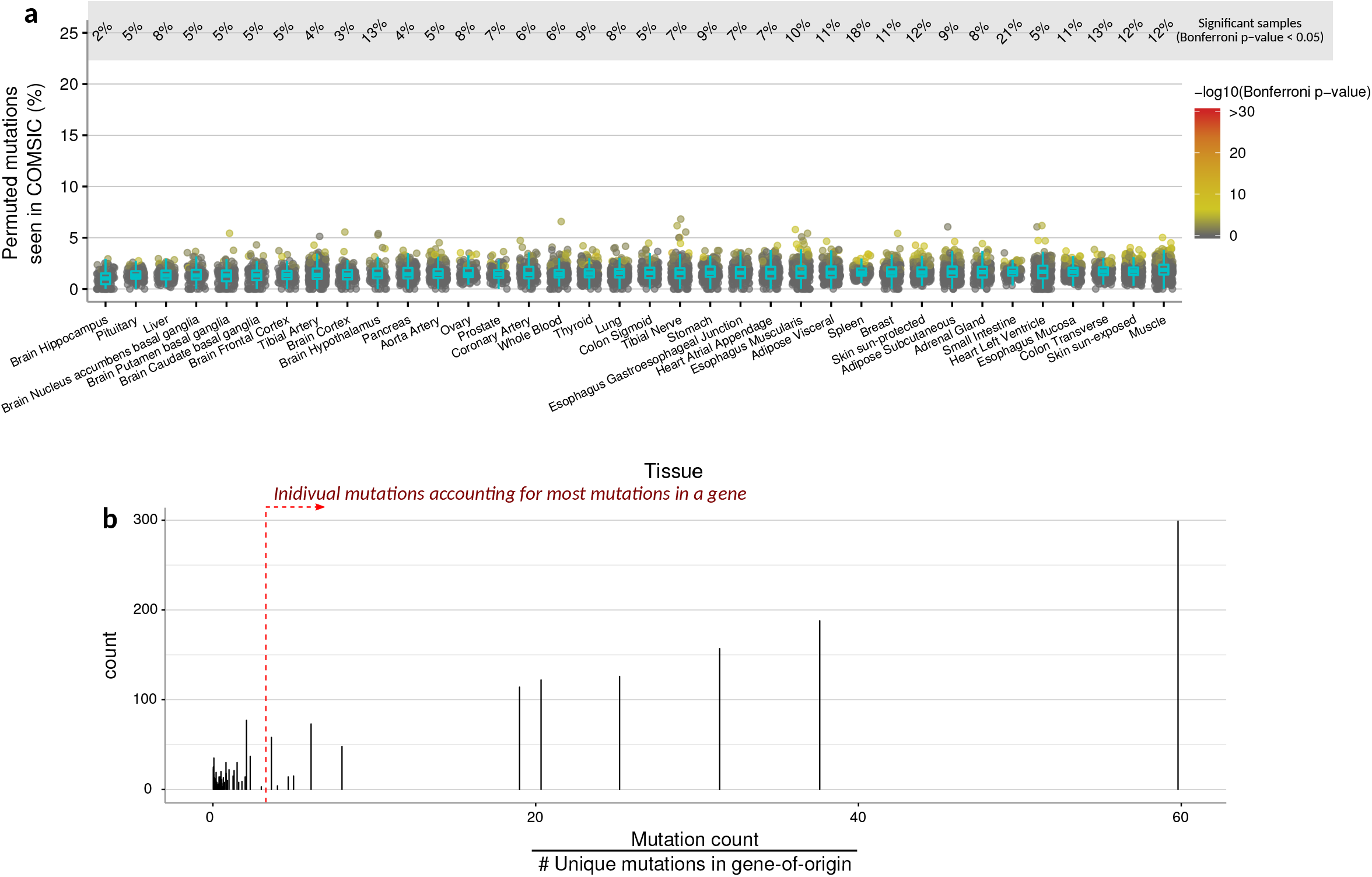
Negative controls and filters for cancer mutation enrichment in non-disease human tissues. **a,** Percentage of randomly permuted mutations (see Methods) in non-disease tissues that overlap with cancer mutation sites (COSMIC) but where the mutation type disagrees (i.e C>A in one dataset and C>G in the other); p-values for enrichment were calculated using a hypergeometric test accounting for sequencing coverage, total number of mutations per sample, and total number of COSMIC mutations, and the three possible alternate alleles that any given reference allele can have (see Methods). P-values are Bonferroni-corrected across all samples. FDR is based on the Benjamini-Hochberg method across all samples. **b,** Histogram of t values (x axis, described in methods) for each mutation in a panel of 31 cancer driver genes. Mutations with high t values account for most of the unique mutated sites in their gene-of-origin, leading to a low diversity in mutations of a given gene. These mutations are likely systematic artifacts and can bias dN/dS ratios^9^.

## METHODS

### Raw RNA-seq retrieval and processing

We downloaded raw RNA sequencing reads from the dbGAP GTEx^23^ project version 7 (phs000424.v7.p2). We included 36 tissues with a total of 7,584 samples (Supp. Table 2) after applying all filters in the mutation calling pipeline (see below). We also processed DNA-exome sequencing reads from 105 whole-blood samples that had a matched RNA-seq sample.

Reads were mapped to the human genome Hg19 (NCBI build GRCh37.p13) using STAR^46^ with the following parameters: requiring uniquely mapping reads (--outFilterMultimapNmax 1), clipping 6 bases in the 5’ end of reads (--clip5pNbases 6), keeping reads with 10 or fewer mismatches and less than 10% mismatches of the read length that effectively mapped to genome (--outFilterMismatchNmax 10, --outFilterMismatchNoverLmax 0.1). To avoid germline variant contamination during the somatic mutation calling phase, all SNPs from dbSNP^47^ v142 were masked to Ns and ignored in downstream processing (we used SNPs annotated in https://ftp.ncbi.nlm.nih.gov/snp/organisms/human_9606_b142_GRCh37p13/BED).

After mapping we removed duplicate reads to avoid biases arising from PCR duplicates during library preparation using a custom python script.

### DNA somatic mutation calling from RNA-sequencing

Identifying DNA variants from RNA-seq requires filtering out many sources of false positives. These include sequencing errors, RNA editing events, mapping errors around splice junctions, germline variants, and other sequencing/mapping biases. To address these issues, we developed a custom mutation calling pipeline that borrows ideas from classical DNA-based variant calling coupled with extra filters that to increase the fidelity of the mutation calls from RNA-seq.

After mapping the raw RNA-seq reads, the pipeline consists of three main parts: 1) identifying positions with two base calls, 2) removing germline variants, and 3) filtering out other potentially spurious variants.

#### Identifying positions with two base calls

After mapping, bam files were scanned to identify genomic positions that were covered by reads having exactly two different base calls. Given the intrinsic sequencing error rate, we only considered positions with high coverage and high sequencing quality for this step. Stringent cutoffs were set for coverage (c >= 40 reads) and sequencing quality (q_s_ >=30 in Phred scale); this is in comparison twice what some DNA-based calling algorithms have used^21^. Finally, only positions in which the minor allele was supported by at least 6 reads were considered (n >= 6). In combination these parameters define a theoretical distribution of the probability of observing a mutation due to sequencing errors, which is small for a wide range of sequence coverage levels (Supp. Fig. 1a). In addition we included a filter to only consider variants with a probability of sequencing error p<0.0001 (see filters below). These strict cutoffs ensure a low probability of observing sequencing errors even in highly covered genes; nonetheless we still apply further filters to account for sequencing errors (see below).

#### Identifying and eliminating germline variants

Common and rare germline variants have to be excluded for the proper identification of somatic mutations. To eliminate common variants a strict filter was applied by using a human genome masked with Ns for positions known to have common variants from dbSNP v142 (see above).

To eliminate all other germline variants, including rare ones, we utilized the low confidence germline variants called by GTEx. These calls were made by GTEx using GATK’s HaplotypeCaller v3.4 using whole-genome sequencing data at 30x coverage from whole blood samples. We specifically used the vcf file *GTEx_Analysis_2016-01-15_v7_WholeGenomeSeq_652Ind_GATK_HaplotypeCaller.vcf* which contains all germline variants before filtering for MAF and low-quality variants. Our goal was to exclude as many germline variants as possible, so we reasoned that using all germline variants called by GTEx – including the low-quality ones – was the safest option to minimize germline mutation contamination in our somatic mutation calls. While these variants were called in whole-blood samples, germline variants should be present in all tissues of an individual and as such these variants were excluded in a per-individual basis across all tissues of that individual. Only sites that had heterozygous or homozygous variants for the alternate allele were excluded from the mutation calls of the given individual.

#### Filtering out artifacts

A total of 13 filters were applied to exclude a variety of artifacts:

1. Blacklisted regions. We excluded sex chromosomes, unfinished chromosomal scaffolds, the mitochondrial genome and the HLA locus in chromosome 6 which is known to contain a high density of germline variants^48^ making accurate mapping challenging, and is hence a potential source of false-positive calls.
2. RNA edits. The most prevalent type of RNA-editing in humans is Adenine-to-Inosine (A>I) which is observed as an A>G/T>C substitution in sequencing data. The Rigorously Annotated Database of A-to-I RNA editing (RADAR)^49^ has extensively curated RNA-edit events including calls identified using the GTEx data^50^. We also obtained RNA-edit information from the Database of RNA-editing (DARNED) that includes further edit types curated from published studies^51^. We excluded all positions described in RADAR v2 (http://lilab.stanford.edu/GokulR/database/Human_AG_all_hg19_v2.txt) and in DARNED (https://darned.ucc.ie/static/downloads/hg19.txt), and observed an average decrease of 10% mutations per sample in our RNA-sequencing calls but not in our DNA-sequencing calls from GTEx (Supp. Fig. 2a), indicating that we are indeed eliminating real RNA-edit events that would otherwise be false-positive mutation calls.
3. Splice junction artifacts. Splice junctions are difficult to resolve during mapping because a gap has to be introduced in reads spanning a splice junction to map it to the corresponding exons in the genome. We observed that the mutation rate was higher close to annotated exon ends and it stabilized at ~7 bp away from the exon end across all tissues (Genecode v19 genes v7 annotation file) (Supp. Fig. 1b). Most of these are likely mapping artifacts and we therefore excluded all mutations located less than 7 bp away from an annotated exon end.
4. Sequencing errors. While sequencing errors are unlikely to be found given the parameters established in the first part of the mutation calling pipeline (see above), we additionally filtered out mutations that had a probability of being sequencing errors greater than 0.01%. This probability was calculated using the upper tail integral of the binomial where the number of successes is the number of reads supporting the alternate allele, the number of events is the coverage in that position, and the probability of success is the conservative assumption of p=0.001 which equals our cutoff of phred score 30 during the first part of the pipeline (see above). This is extremely conservative because it does not incorporate the probability of observing the exact same base call across all reads supporting the alternate allele.
5. Read position bias. To eliminate any systematic bias of a mutation being consistently called around the same position along reads supporting it versus the rest of the reads, we excluded mutations that had a p-value less than 0.05 when applying a Mann-Whitney U test of the positions in the read supporting the alternate allele vs the positions supporting the reference allele. For these tests we used BCFtools^52^ *mpileup*.
6. Mapping quality bias. We excluded mutations that had a p-value less than 0.05 when applying a Mann-Whitney U test comparing mapping quality scores of the base calls supporting the alternate allele vs the mapping quality scores of reads supporting the reference allele. For these tests we used BCFtools^52^ *mpileup*.
7. Sequence quality bias. We excluded mutations with a p-value less than 0.05 when applying a Mann-Whitney U test comparing sequencing quality scores of base calls supporting the alternate allele vs the scores of base calls supporting the reference allele. For these tests we used BCFtools^52^ *mpileup*.
8. Strand bias. We excluded mutations with a p-value less than 0.05 when applying a Mann-Whitney U test comparing strand bias of bases supporting the reference and alternate allele (i.e., cases where mutations were only observed on one strand were excluded; this is not related to the strand asymmetry we observed for some mutation types). For these tests we used the ‘Mann-Whitney U test of Mapping Quality vs Strand Bias’ of BCFtools^52^ *mpileup*.
9. Variant distance bias. We excluded variants that showed a high or low mean pairwise distance between the alternate allele positions in the reads supporting it. Similar to the *read position bias* filter, this ensures that we filter mutations that are consistently observed around the same region of all the reads that support it. We used a cut off of p<0.05 for a two-tail distribution of simulated mean pairwise distances from the BCFtools^52^ implementation.
10. RNA-specific allele frequency bias. We observed an enrichment of variants having a variant allele frequency (VAF) greater than 0.9 only in mutation calls from RNA-sequencing but not from the matched DNA-sequencing samples. This RNA-specific bias could be due to several factors, including allele specific expression leading to enrichment of the alternate allele, RNA editing, or systematic artifacts during RNA extraction, library construction, and sequencing. We took a conservative approach and excluded all mutations that had a VAF greater than 0.7.
11. Tissue-specific mutation effects. To eliminate false-positives arising by systematic artifacts of unknown origin, we first looked for recurrent mutations observed in many samples of a given tissue. While these mutations could be real and have biological impact, they may also reflect a shared systematic artifact that is producing the same exact mutation across several samples of the same tissue. We decided to take a conservative approach and eliminated all mutations that were called in at least 40% of the samples in one tissue. Even though we labelled these mutations on a per-tissue basis, once identified we removed them from any sample in any tissue that had them.
12. Overall systematic mutation bias. We further eliminated mutations that were present in at least 4% of all samples. Similarly, as in the previous step these mutations are more likely to have originated from a systematic artifact.
13. Hyper-mutated samples. We excluded samples that had an excess of mutations compared to what it was expected from sequencing depth and biological factors. To do so we looked at the residuals after applying a linear regression on mutation numbers using as features sequencing depth, age, sex, and BMI. We observed 48 samples that had residual values greater than 1500 (i.e. they had >1500 more mutations than expected by other factors) (Supp. Fig. 1e) and excluded them from further analysis, leaving 7,584 remaining. We did not observe any hypo-mutated samples having similar residual values in the opposite direction.

### Method validation

To validate the precision of our DNA somatic mutation calls from RNA-seq, we sought to compare those calls to DNA sequencing from the exome. The GTEx Consortium performed DNA exome capture followed by sequencing in blood samples with a median coverage of 80x.

From the GTEx dbGaP repository (phs000424.v7.p2) we downloaded raw DNA exome sequencing runs from 105 randomly chosen donors. We mapped the reads and called mutations identically to our RNA-seq samples. Throughout the study we used the mutation calls from exome to validate certain results when appropriate.

To validate the overall mutation calling pipeline we matched RNA-seq to exome DNA-seq samples of the same individuals. Then, we asked what percentage of the mutation calls from RNA-seq had reads supporting the alternate allele in the matched exome DNA-seq. To account for coverage differences between RNA-seq and DNA-seq, we focused this analysis only on mutation calls for which we had reasonable power of detection to validate mutations in DNA-seq data. This is especially relevant because in highly expressed genes, sequencing coverage can be >1000x in RNA-seq; therefore we could detect mutations with VAF as low as 0.006 for a 1000x-covered position (given our minimum alternate allele 6-read cutoff described above). In comparison, in exome DNA-seq given the 80x median coverage we expect to see only 0.48 reads supporting the alternate allele of a variant with VAF = 0.006. We therefore devised a way to account for this coverage difference between RNA-seq and DNA-seq, and only compared positions where we had power of detection in both experiments. We first made the following calculation:

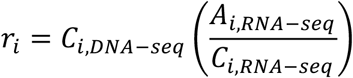

Where *C_i_* and *A_i_* are total and alternate read counts for a mutation in position *i; r_i_* effectively represents the number of expected reads to support the alternate allele in DNA-seq at position *i* given the VAF observed in RNA-seq. As shown in Supplementary Figure 1d, we took a conservative approach and only compared genomic positions where *r* ≥ 8, to ensure sufficient power of validation.

For all positions selected we then calculated the percentage of mutations observed in an RNA-seq sample that had at least one read supporting the variant allele in the exome DNA-seq, effectively resulting in a false-discovery rate. When assigning a random alternate allele (not including the reference allele among the choices) to the mutated positions in RNA-seq we did not observe support for >98% of the position-permuted mutations, thus proving the validity of this FDR calculation (Supp. Fig. 1c).

### Discovery of mutation associations with biological and non-biological factors

We first sought to identify and subtract the effects of non-biological factors influencing the number of mutations observed per sample. To this end we grouped samples by tissue and performed a linear regression on the mutation numbers using non-biological factors as explanatory variables, which included sequencing depth, transcriptome Shannon diversity, time spent in the PAXgene fixative (SMTSPAX from GTEx sample attributes), total ischemic time (SMTSISCH from GTEx sample attributes), and RNA integrity number (SMRIN from GTEx sample attributes). We assessed the significance of association between mutation number and any factor by the p-value of each coefficient in a multiple regression (Supp. Table 4).

To discover biological factors associated with mutation numbers, we took the residuals of the previous linear regression (constructed with non-biological explanatory variables) and performed another regression using biological factors as explanatory variables. Biological factors included: biological sex, age, BMI, and the first three genotype principal components constructed from the ~10^7^ SNPs with MAF >= 0.1 available from GTEx. Significance was assessed by the p-value of each coefficient in a multiple regression (Supp. Table 4). Additionally, Spearman correlations were independently performed between each biological factor and the residuals from the linear regression between mutation number and non-biological factors.

### tSNE

tSNE was performed on a matrix of *m* rows and *n* columns, where *m* are samples and *n* are mutation type counts normalized by their corresponding sequence content in the sample transcriptome. We used a 1536-type mutation profile including two base-pairs upstream and downstream of the mutation site (pentanucleotide profile). The normalized mutation count (*mut_i_*) by sequence content in the transcriptome was obtained as follows:

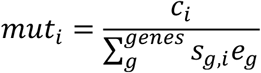

Where *c_i_* is the count of the mutation type *i, s_g,i_* is the number of occurrences of the *i* sequence context in gene *g, e_g_* is the expression in TMP of gene *g*, and *genes* are all genes with TPM >= 1 in the given sample. This calculation was performed separately for every sample. We used the tSNE implementation in R (Rtsne) with parameters dims = 2, max_iter = 500, perplexity = 30, pca = TRUE, theta = 0.5.

To quantify the clustering among different groupings in the tSNE two-dimensional space, we calculated a silhouette score (SS) for each group defined in Figure 2f by first obtaining a silhouette score (*s_i_*) for each sample in each group:

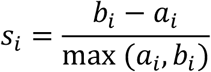

Where *a_i_* is the average distance of sample *i* to all other samples inside the group, and *b_i_* is the average distance of sample *i* to samples outside the group. We then calculated the average score *s* of all samples in a given group and the 95% confidence intervals based on bootstrapping 10,000 times.

We performed the individual-based grouping by calculating silhouette scores for all tissues from 20 randomly selected individuals and then averaging them. The random expectation was calculated by permuting the tissue labels across samples 10 times, repeating the SS calculation across tissues and then averaging all SSs.

### Cell type decomposition for blood and lung samples

We used CIBERSORT^45^ to identify the cell type composition of each whole blood sample in the GTEx data. Briefly, CIBERSORT applies a support vector regression on a gene expression profile(s) using reference gene expression signatures from different cell types, and then retrieves the cell type composition from the signatures in the original expression profile(s). We used the online portal (https://cibersort.stanford.edu/) and the default LM22 expression signatures composed of the most prevalent immune cell types. CIBERSORT was run on the blood gene expression profiles with default parameters: 100 permutations and “absolute” mode.

### Gene expression associations

To avoid population-based effects for all expression associations analyses in this study we only used self-reported Caucasian people.

For each tissue we used individual gene expression to model mutation counts in a linear regression as follows:

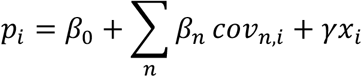

Where *x* is the expression of a given gene in TPM, *cov* is a covariate variable and there are *n* covariates. To estimate the effects of a SNP on the phenotype, γ is calculated for each gene in the genome that has a TPM > 1 in at least 20% of the samples. Significance is measured based on the p-value of a non-zero t-test performed on γ. P-values are adjusted using Bonferroni correction.

When *p_i_* is the vector of mutation loads, we defined it as the residuals after regressing non-biological factors as described previously (see above in “Discovery of mutation associations with biological and non-biological factors”). The covariates included in this model are the first 3 PCs of the whole-genome genotypes, age, sex, and BMI.

In all cases both *p*_i_ and γ were normalized by converting the values into quantiles and mapping them to the corresponding values of the standard normal distribution quantiles, following standard GTEx practices^23^.

### Gene Ontology analysis

For gene expression associations with mutation load, we assessed enrichment in GO biological processes using GOrilla^53^. We used as input a ranked list of genes based on the number of tissues they were significant in (and only including genes that were tested in more than the median number of tissues all genes were tested in, p<0.05 after Bonferroni correction, see above in “Gene expression associations”) and breaking ties by significance. We then used REVIGO^54^ to obtain non-redundant categories.

### Expression associations with genomic instability genes

We selected a panel of genes known to be involved in different pathways of DNA repair or translesion replication (Fig. 4) and performed association analyses between their expression and the mutational load for all different mutation types on a per-tissue basis. We followed the same linear regression strategy as described above in “Gene expression associations”. We devised customized strategies to calculate FDRs for individual gene-tissue associations, gene enrichment across all tissues, and pathway (gene group) associations in each tissue.

For individual gene-tissue associations, we calculated an FDR based on the distribution of p-values of the linear regressions from the tests of all genes in the genome (see above for additional filters). Then, for each gene of interest we calculated the FDR as the percentage of genes from all tests that had an equal or lower p-value.

To obtain an FDR of the enrichment for different pathways across tissues, for each pathway of interest we obtained the p-values from the individual linear regressions of each gene. We then created 10,000 groups with randomly selected genes with the same size as the pathway of interest and within the tissue of interest, and performed linear regressions with mutation load (as describe above in “Gene expression associations”). Finally, for each p-value in the original group we calculated the FDR by dividing the percentage of p-values of equal or lower value in all permuted regressions by the percentage of p-values of equal or lower value in the original regressions. For example, if 0.1% of permuted regressions reached p<0.001, compared to 1% of unpermuted regressions, the FDR at p<0.001 would be 0.1/1 = 10%.

Lastly, to calculate FDR for the enrichment of individual genes across all tissues we used a similar strategy as the FDR calculation for gene pathways described in the previous paragraph. We first obtained the p-values of individual linear regression of a given gene across all tissues. We then created a permuted set of p-values from the regressions of one gene from each of the 10,000 permuted groups in all tissues. Finally, we calculated the FDR of the p-values in the original group (one across all tissues) by dividing the percentage of p-values of equal or lower p-value in all permuted regressions by the percentage of p-values of equal or lower value in the original regressions.

### Chromatin analysis

To address the influence of chromatin on somatic mutations, we first manually mapped tissues from GTEx to tissues of the Roadmap Epigenomics Project^28^. We were able to do such mapping for 18 tissues (Supp. Table 7). For each tissue, on a per-exon basis we calculated both mutation rates and signal for H3K36me3, H3K4me1, H3K4me3, H3K27me3, H3K9me3.

To account for sequencing depth (expression) of an exon when calculating mutation rates, we assigned exons to 100 bins based on their sequencing depth, where each bin contains 1% of exons. We then calculated the median number of mutations observed in each bin, and finally the mutation rate per exon was obtained by subtracting expected number of mutations (the median for all exons in that bin) from the observed number for that exon.

Chromatin signal for each exon was obtained from Roadmap ChIP-seq data. We calculated the average base-pair ratios of IP/input obtained from the Roadmap bigwig files.

Significance of the association between each chromatin mark and mutation rate was assessed by applying a linear regression on the mutation rate using all chromatin marks as features.

### Mutation enrichment in COSMIC cancer mutations

We downloaded the entire set of cancer mutations from COSMIC^36^ v86 and we further filtered them to only keep single nucleotide variants without indels. For each sample we calculated the percentage (overlap) of their mutations that are present in COSMIC mutations, and we calculated the significance of the overlap using the integral of the upper tail from a hypergeometric distribution, that is p(X > k), where X follows a hypergeometric distribution with parameters *K* (number of mutations in sample), *n* (number of COSMIC mutations whose positions were covered by >= 40 reads in the RNA-seq sample), *N × 3* (the total number of base pairs covered by >= 40 reads in the RNA-seq sample multiplied by the three possible alternate alleles that any reference can have), and *k* (the overlap of mutations, *K* and *n*). As a control, for each sample this calculation was repeated using a permuted set of mutations. This set was constructed by randomly selecting genomic positions that were covered by >=40 reads in the sample and then mutations were simulated in those positions, importantly we conserved the number of reference alleles and their corresponding alternate alleles from the original mutations in this permuted set.repeated.

### dN/dS analysis

To calculate dN/dS ratios we applied a previously described method^38^ that uses a Poisson distribution to model the number of mutations with different impacts (i.e. synonymous vs non-synonymous). Briefly, the Poisson distribution is based on the relative content of a mutation type (e.g. C>T) across all types, the total content of that mutation type, and the density of mutations per site. For non-synonymous mutations an extra parameter represents the effect of selection (dN/dS), and maximum-likelihood estimates are calculated by Poisson regression for all parameters. This framework accounts for different substitution rates across different genes as well as sequence composition. We used dndsloc, which is the implementation of this method in R^9^ (https://github.com/im3sanger/dndscv).

### Mutation analyses on cancer driver genes

We downloaded the list of genes known to contain at least one cancer driver mutation, based on the latest TCGA publication on cancer driver genes^37^(Supp. Table 10). Since we did more in depth analysis in this set of genes, we further removed potential false-positive mutation calls that could bias gene-level analysis but are not necessarily an issue for genome-wide analysis. For some of these genes we observed a high number of total mutations but low number of unique mutations, in other words some mutations accounted for most of the unique mutated sites, which has been previously described as an artifact^9^. To flag these events, for each mutation we calculated a metric *t* as the ratio of the counts of that mutation divided by the unique number of mutations found in the gene-of-origin. Mutations with high *t* values account for most of the unique mutated sites in their gene-of-origin, leading to a low diversity in mutations of a given gene. These mutations may be artifacts and can bias dN/dS ratios^9^. The distribution of *t* values for mutations in this set of genes was bimodal (Supp. Fig. 11b) and we therefore excluded mutations with *r >* 3.5.

To assess the mutation load on these genes and their significance, for each tissue we calculated the mutation rate of these genes, and then calculated whether this mutation rate was higher or lower than expected from the overall mutation rate on these genes across all tissues (Fig. 5b). To do so we calculated the overall mutation rate in cancer driver genes across all tissues (*k*) and then for each tissue we calculated the probability of the number of observed mutations (*n*) in these genes using the binomial distribution [*X* ~ *binom(n,k)*]. For tissues showing more mutations than expected we used the integral of the right tail of the binomial distribution to calculate the probability of observing n mutations, and conversely for tissues showing fewer mutations we used the integral of the left tail of the distribution. FDR was calculated using Benjamini-Hochberg method on the p-values.

We annotated the oncogenic status for the mutations in these driver genes using the oncokb^39^ tool *MafAnnotator.py*.

## Acknowledgements

We would like to thank J. Pritchard, B. Li, and members of the Fraser Lab for helpful feedback. PEGN is supported by a Bio-X Bowes Graduate Student Fellowship. This work was supported by NIH grant 2R01GM097171-05A1.

## REFERENCES

1. Kennedy, S. R., Loeb, L. A. & Herr, A. J. Somatic mutations in aging, cancer and neurodegeneration. Mech. Ageing Dev. 133, 118–126 (2012).

2. Bailey, M. H. et al. Comprehensive Characterization of Cancer Driver Genes and Mutations. Cell 173, 371–385.e18 (2018).

3. Alexandrov, L. B. et al. Signatures of mutational processes in human cancer. Nature 500, 415–421 (2013).

4. Schuster-Bockler, B. & Lehner, B. Chromatin organization is a major influence on regional mutation rates in human cancer cells. Nature 488, 504–507 (2012).

5. Paz Polak, Rosa Karlic, Amnon Koren, Robert Thurman, Richard Sandstrom, Michael S. Lawrence, A. R. & Eric Rynes, Kristian Vlahoviček, J. A. S. & S. R. S. Cell-of-origin chromatin organization shapes the mutational landscape of cancer. Nature 518, 360–364 (2015).

6. García-Nieto, P. E. et al. Carcinogen susceptibility is regulated by genome architecture and predicts cancer mutagenesis. EMBO J. 36, 2829–2843 (2017).

7. Smith, K. S., Liu, L. L., Ganesan, S., Michor, F. & De, S. Nuclear topology modulates the mutational landscapes of cancer genomes. Nat. Struct. Mol. Biol. 24, 1000–1006 (2017).

8. Yadav, V. K., DeGregori, J. & De, S. The landscape of somatic mutations in protein coding genes in apparently benign human tissues carries signatures of relaxed purifying selection. Nucleic Acids Res. 44, 2075–2084 (2016).

9. Martincorena, I. et al. Universal Patterns of Selection in Cancer and Somatic Tissues. Cell 171, 1029–1041.e21 (2017).

10. Temko, D., Tomlinson, I. P. M., Severini, S., Schuster-Böckler, B. & Graham, T. A. The effects of mutational processes and selection on driver mutations across cancer types. Nat. Commun. 9, 1857 (2018).

11. Zapata, L. et al. Negative selection in tumor genome evolution acts on essential cellular functions and the immunopeptidome. Genome Biol. 19, 67 (2018).

12. Samstein, R. M. et al. Tumor mutational load predicts survival after immunotherapy across multiple cancer types. Nat. Genet. (2019). doi:10.1038/s41588-018-0312-8

13. Bose, S., Deininger, M., Gora-Tybor, J., Goldman, J. M. & Melo, J. V. The presence of typical and atypical BCR-ABL fusion genes in leukocytes of normal individuals: biologic significance and implications for the assessment of minimal residual disease. Blood 92, 3362–7 (1998).

14. Xie, M. et al. Age-related mutations associated with clonal hematopoietic expansion and malignancies. Nat. Med. 20, 1472–1478 (2014).

15. Martincorena, I. et al. High burden and pervasive positive selection of somatic mutations in normal human skin. Science (80-.). 348, 880–886 (2015).

16. Bae, T. et al. Different mutational rates and mechanisms in human cells at pregastrulation and neurogenesis. Science (80-.). eaan8690 (2017). doi:10.1126/science.aan8690

17. Lodato, M. A. et al. Aging and neurodegeneration are associated with increased mutations in single human neurons. Science (80-.). eaao4426 (2017). doi:10.1126/science.aao4426

18. Martincorena, I. et al. Somatic mutant clones colonize the human esophagus with age. Science (80-.). eaau3879 (2018). doi:10.1126/SCIENCE.AAU3879

19. Yokoyama, A. et al. Age-related remodelling of oesophageal epithelia by mutated cancer drivers. Nature 565, 1 (2019).

20. Lee-Six, H. et al. The landscape of somatic mutation in normal colorectal epithelial cells. bioRxiv 416800 (2018). doi:10.1101/416800

21. Xu, C. A review of somatic single nucleotide variant calling algorithms for next-generation sequencing data. Comput. Struct. Biotechnol. J. 16, 15–24 (2018).

22. Enge, M. et al. Single-Cell Analysis of Human Pancreas Reveals Transcriptional Signatures of Aging and Somatic Mutation Patterns. Cell 171, 321–330 (2017).

23. Consortium, G. Genetic effects on gene expression across human tissues. (2017). doi:10.1038/nature24277

24. Leija-Salazar, M., Piette, C. & Proukakis, C. Review: Somatic mutations in neurodegeneration. Neuropathol. Appl. Neurobiol. 44, 267–285 (2018).

25. Lindeboom, R. G. H., Supek, F. & Lehner, B. The rules and impact of nonsense-mediated mRNA decay in human cancers. Nat. Genet. 48, 1112–1118 (2016).

26. Igo Martincorena, I., Raine, K. M., Davies, H., Stratton, M. R. & Campbell, P. J. Universal Patterns of Selection in Cancer and Somatic Tissues. (2017). doi:10.1016/j.cell.2017.09.042

27. Van Der Maaten, L. & Hinton, G. Visualizing Data using t-SNE. Journal of Machine Learning Research 9, (2008).

28. Kundaje, A. et al. Integrative analysis of 111 reference human epigenomes. Nature 518, 317–330 (2015).

29. Haradhvala, N. J. J. et al. Mutational Strand Asymmetries in Cancer Genomes Reveal Mechanisms of DNA Damage and Repair. Cell 164, 538–549 (2016).

30. Kucab, J. E. et al. A Compendium of Mutational Signatures of Environmental Agents. Cell 0, (2019).

31. Sharma, S. et al. Mitochondrial hypoxic stress induces widespread RNA editing by APOBEC3G in natural killer cells. Genome Biol. 20, 1–17 (2019).

32. Kim, H. & Kim, Y.-M. Pan-cancer analysis of somatic mutations and transcriptomes reveals common functional gene clusters shared by multiple cancer types. Sci. Rep. 8, 6041 (2018).

33. Eliopoulos, A. G., Havaki, S. & Gorgoulis, V. G. DNA Damage Response and Autophagy: A Meaningful Partnership. Front. Genet. 7, 204 (2016).

34. Wang, Y. JimMY on the stage: Linking DNA damage with cell adhesion and motility. Cell Adh. Migr. 4, 166–8

35. Richman, S. Deficient mismatch repair: Read all about it (Review). Int. J. Oncol. 47, 1189–202 (2015).

36. Tate, J. G. et al. COSMIC: the Catalogue Of Somatic Mutations In Cancer. Nucleic Acids Res. 47, 941–947 (2018).

37. Bailey, M. H. et al. Comprehensive Characterization of Cancer Driver Genes and Mutations. Cell 173, 371–385.e18 (2018).

38. Wong, C. C. et al. Inactivating CUX1 mutations promote tumorigenesis. Nat. Genet. 46, 33–38 (2014).

39. Chakravarty, D. et al. OncoKB: A Precision Oncology Knowledge Base. JCO Precis. Oncol. 1, 1–16 (2017).

40. Martincorena, I. & Campbell, P. J. Somatic mutation in cancer and normal cells. Science (80-.). 349, 1483–1489 (2015).

41. Cancer Genome Atlas Research Network, T. et al. Comprehensive and Integrated Genomic Characterization of Adult Soft Tissue Sarcomas. (2017). doi:10.1016/j.cell.2017.10.014

42. Chang, M. T. et al. Accelerating Discovery of Functional Mutant Alleles in Cancer. Cancer Discov. 8, 174–183 (2018).

43. Gao, J. et al. 3D clusters of somatic mutations in cancer reveal numerous rare mutations as functional targets. Genome Med. 9, 4 (2017).

44. Tomasetti, C. & Vogelstein, B. Cancer etiology. Variation in cancer risk among tissues can be explained by the number of stem cell divisions. Science 347, 78–81 (2015).

45. Newman, A. M. et al. Robust enumeration of cell subsets from tissue expression profiles. Nat. Methods 12, 453–457 (2015).

46. Dobin, A. et al. STAR: ultrafast universal RNA-seq aligner. Bioinformatics 29, 15–21 (2013).

47. Sherry, S. T. et al. dbSNP: the NCBI database of genetic variation. Nucleic Acids Res. 29, 308–311 (2001).

48. Buhler, S. & Sanchez-Mazas, A. HLA DNA sequence variation among human populations: molecular signatures of demographic and selective events. PLoS One 6, e14643 (2011).

49. Ramaswami, G. & Li, J. B. RADAR: a rigorously annotated database of A-to-I RNA editing. Nucleic Acids Res. 42, D109–D113 (2014).

50. Tan, M. H. et al. Dynamic landscape and regulation of RNA editing in mammals. Nature 550, 249–254 (2017).

51. Kiran, A. M., O’Mahony, J. J., Sanjeev, K. & Baranov, P. V. Darned in 2013: inclusion of model organisms and linking with Wikipedia. Nucleic Acids Res. 41, D258–D261 (2012).

52. Li, H. et al. The Sequence Alignment/Map format and SAMtools. Bioinformatics 25, 2078–2079 (2009).

53. Eden, E., Navon, R., Steinfeld, I., Lipson, D. & Yakhini, Z. GOrilla: a tool for discovery and visualization of enriched GO terms in ranked gene lists. BMC Bioinformatics 10, 48 (2009).

54. Supek, F., Bošnjak, M., Škunca, N. & Šmuc, T. REVIGO Summarizes and Visualizes Long Lists of Gene Ontology Terms. PLoS One 6, e21800 (2011).

