## Supplementary notes for "The somatic mutation landscape of the human body"

### Supplementary Note 1. Somatic mutation burden of human tissues is partially associated with number of stem cell divisions in the tissue

Tomasetti and Vogelstein[1] found a strong correlation between lifetime risk of many cancer types and the total number of stem cell divisions within the tissue of origin likely originating each cancer. They proposed a mechanism by which the number of mutations a cell experiences linearly increases with the number of cell divisions, and the greater the number of divisions the more likely carcinogenic mutations will appear and consequently lead to cancer. To address this hypothesis we matched GTEx tissues to cancer types used in Tomasetti and Vogelstein, 2015[1]. Matching was done only for cancer types whose cell-of-origin corresponds to the most common cell type in a GTEx tissue (Supp. Table 13). Across different mutation types we observed positive but mildly significant correlations between mutation load of GTEx tissues and number of stem cell divisions of its matching tissue from Tomasetti and Vogelstein (Supp. Fig. 5). C>G mutations show the highest correlation (spearman $\rho = 0.52$, $p = 0.098$), followed by T>A (spearman $\rho = 0.48$, $p = 0.14$), C>T (spearman $\rho = 0.4$, $p = 0.23$), C>A (spearman $\rho = 0.36$, $p = 0.28$), T>G (spearman $\rho = 0.15$, $p = 0.67$), and T>C (spearman $\rho = 0.018$ $p = 0.96$). Small intestine and pancreas are the two outlier tissues, with small intestine showing greater number of mutations than expected by stem cell divisions and pancreas lower than expected. These results suggest that the number of stem cell divisions contributes to mutation load in most tissues.

### Supplementary Note 2. Individual tissue-gene associations between global mutation load and expression of DNA repair genes

We found 14 tissue-gene associations at a maximum FDR of 20% (Fig 4c, blue asterisks). These associations include *OGG1* expression with higher mutation load in ovary (Supp. Fig. 8c; $p = 5 \times 10^{-7}$, FDR = 0.002), *NEIL1* expression association with higher mutation load in liver (Supp. Fig. 8d; $p = 2.95 \times 10^{-12}$, FDR = 0.02), and *MSH3* expression association with lower mutation load in the adrenal gland (Supp. Fig. 8f; $p = 0.0001$, FDR = 0.003). P-values in this section are from a linear regression applied on mutation counts using gene expression as well as other biological and technical covariates (see methods).

These associations may or may not reflect causal relationships between these genes and somatic mutation rates. Evidence that they may not be causal comes from previous studies that have observed the opposite direction of relationship. For example, the expression of *NEIL1* has been shown to negatively correlate with mutation load in several cancers[2] and a knockout in mouse liver leads to a metabolic disorder thought to be caused by an increase in DNA damage[3]; similarly *OGG1* knockout in mouse leads to a predisposition of ovarian cancer[4]. In comparison, *MLH1* has been observed to increase cancer risk in lung cancer[5], which suggests a potential causal link between *MLH1* gene expression and mutation load in lung (Supp. Fig. 8e)

### Supplementary Note 3. Evidence for involvement in mutagenesis of DNA repair genes associated with mutation load across several tissues of the human body

*XPA*, *XPC* and *DDB1* knockouts in human cell lines[6,7] and mouse[8] have been shown to cause increased mutagenesis. Additionally, there is evidence that inactivation of both *MSH6* and *PMS2* lead to an increase in mutagenesis due to deficiencies in MMR[9–11]. Finally, *NEIL2* is known to repair DNA damage in actively transcribed regions of the genome and cells lacking this gene show increased levels of DNA damage[12], and *NEIL2* expression is correlated in the same direction we observed with cancer mutagenesis[2]. These associations are in the same direction as the ones observed between the expression of these genes and mutation load from this study.

**REFERENCES**


1. Tomasetti, C. & Vogelstein, B. Cancer etiology. Variation in cancer risk among tissues can be explained by the number of stem cell divisions. *Science* **347**, 78–81 (2015).
2. Shinmura, K. *et al.* Abnormal Expressions of DNA Glycosylase Genes NEIL1, NEIL2, and NEIL3 Are Associated with Somatic Mutation Loads in Human Cancer. *Oxid. Med. Cell. Longev.* **2016**, 1–10 (2016).
3. Vartanian, V. *et al.* The metabolic syndrome resulting from a knockout of the NEIL1 DNA glycosylase. *Proc. Natl. Acad. Sci.* **103**, (2006).
4. Xie, Y. *et al.* Deficiencies in Mouse Myh and Ogg1 Result in Tumor Predisposition and G to T Mutations in Codon 12 of the K-Ras Oncogene in Lung Tumors. *Cancer Res.* **64**, 3096–3102 (2004).
5. Cooper, W. A. *et al.* Prognostic significance of DNA repair proteins MLH1, MSH2 and MGMT expression in non-small-cell lung cancer and precursor lesions. *Histopathology* **52**, 613–22 (2008).
6. Sassa, A., Kamoshita, N., Kanemaru, Y., Honma, M. & Yasui, M. Xeroderma Pigmentosum Group A Suppresses Mutagenesis Caused by Clustered Oxidative DNA Adducts in the Human Genome. *PLoS One* **10**, e0142218 (2015).
7. Lovejoy, C. A., Lock, K., Yenamandra, A. & Cortez, D. DDB1 maintains genome integrity through regulation of Cdt1. *Mol. Cell. Biol.* **26**, 7977–90 (2006).
8. Wijnhoven, S. W. *et al.* Age-dependent spontaneous mutagenesis in Xpc mice defective in nucleotide excision repair. *Oncogene* **19**, 5034–5037 (2000).
9. van Boxtel, R. *et al.* Lack of DNA mismatch repair protein MSH6 in the rat results in hereditary non-polyposis colorectal cancer-like tumorigenesis. *Carcinogenesis* **29**, 1290–1297 (2008).
10. Andrew, S. E. *et al.* Mutagenesis in PMS2- and MSH2-deficient mice indicates differential protection from transversions and frameshifts. *Carcinogenesis* **21**, 1291–5 (2000).
11. van Oers, J. M. M. *et al.* PMS2 endonuclease activity has distinct biological functions and is essential for genome maintenance. *Proc. Natl. Acad. Sci. U. S. A.* **107**, 13384–9 (2010).
12. Banerjee, D. *et al.* Preferential Repair of Oxidized Base Damage in the Transcribed Genes of Mammalian Cells. *J. Biol. Chem.* **286**, 6006–6016 (2011).
